# Functional MRI signals at and beyond 1 Hz are coupled to brain states and predict spontaneous neural activity

**DOI:** 10.1101/2025.10.13.681720

**Authors:** Leandro P. L. Jacob, Sydney M. Bailes, Stephanie D. Williams, Carsen Stringer, Bruce R. Rosen, Jonathan R. Polimeni, Laura D. Lewis

**Author notes:** Corresponding authors –.

## Abstract

Technological advances have enabled fMRI acquisition with high temporal resolution, enabling brainwide imaging in just a few hundreds of milliseconds. However, the relationship between fast hemodynamic signals and spontaneous neural activity in the resting state is not yet well understood, limiting our ability to infer neural processes from these fast data. We hypothesized that high-frequency fMRI signals are linked to spontaneous neural activity tied to vigilance states, and that these high-frequency signals could be used to infer the dynamic variations in neuronal activity indexed by EEG neural rhythms. Using fast fMRI (TR=378 ms) and simultaneous EEG in 27 humans drifting between sleep and wakefulness, we found that fMRI power increased during NREM sleep (compared to wakefulness) in frequencies up to 1.3 Hz. High-frequency fMRI power was also correlated to canonical arousal-linked EEG rhythms (alpha and delta), with spatiotemporal cross-correlation patterns reflecting both shared arousal dynamics and rhythm-specific signatures. Using machine learning, we found that EEG alpha and delta power can be decoded from high-frequency fMRI signals in subjects held-out from the training set, showing that the high-frequency components of fMRI signals contain neurally-coupled information robust enough to generalize across individuals. These results reveal that high-frequency fMRI signals are coupled to dynamically varying brain states, and that fast fMRI allows for temporally precise quantification of spontaneous neural activity.

**Significance Statement:** Functional MRI enables non-invasive brainwide neuroimaging in humans, but classically had limited temporal resolution. Recently, studies demonstrated that fast fMRI sampling can detect stimulus-evoked neural activity at subsecond timescales. However, it was unknown whether spontaneous neural activity is represented in fast fMRI signals. Using fMRI and simultaneous EEG in subjects who fell asleep, we found that EEG neural activity associated with sleep and wakefulness is represented in high-frequency fMRI signals. The results reveal that fast fMRI sampling can be used to investigate the spontaneous neural activity underlying brain states with high temporal precision.

## Introduction

Functional magnetic resonance imaging (fMRI) allows for non-invasive whole-brain neuroimaging by inferring neural activity via the BOLD (blood oxygenation level dependent) signal. Historically, most fMRI studies acquired data with sampling intervals on the order of seconds, and analyzed the BOLD signal using hemodynamic response functions that assume a sluggish BOLD response to neural events. However, over the past decade fast imaging has become widely available (Setsompop et al., 2016), and this faster imaging can enable detection of neural activity at subsecond timescales during structured tasks (Lewis et al., 2016). Furthermore, fast fMRI signals can enable accurate decoding of stimulus-evoked neural activity from short windows, enabling imaging of neural responses with high temporal precision (Miyawaki et al., 2025; Rafeh et al., 2025).

An open question is whether fast fMRI can be used to detect spontaneous neural activity. Spontaneous neural dynamics are fundamental to the study of brain states, such as alertness or sleep, but are particularly difficult to investigate without the benefits of task structure and trial averaging. Resting-state studies have detected spatially structured BOLD signals in high frequency ranges (Boubela et al., 2013; Chen & Glover, 2015; Gohel & Biswal, 2015; Lee et al., 2013), suggesting that fMRI may display a range of fast responses to spontaneous neural activity. However, it is not known whether high-frequency spontaneous BOLD signals are coupled to specific neural signatures, such as the distinct neural rhythms that vary across vigilance states.

Low-frequency (<0.1 Hz) BOLD signals are strongly coupled to variations in brain states and spontaneous neural rhythms. fMRI power below 0.1 Hz increases during sleep (B. Davis et al., 2016; Fukunaga et al., 2006; Horovitz et al., 2008), and fMRI oscillations below 0.2 Hz can track canonical sleep oscillations measured with simultaneous EEG (Song et al., 2022). However, since prior studies sampled fMRI data on timescales of seconds, they could not resolve faster signals. Fast fMRI could particularly benefit investigations of oscillatory sleep dynamics, which include important activity patterns in the 0.5–4 Hz range (Adamantidis et al., 2019), as this fast sampling could allow for the acquisition of high-frequency BOLD signals that overlap with the actual oscillatory range of this neural activity—possibly even allowing for this activity to be decoded directly from high-frequency BOLD.

Here, we use simultaneous EEG and fast fMRI (TR=378 ms) to determine how fMRI dynamics up to 1.3 Hz change across brain states, and identify their link with canonical arousal-related EEG neural rhythms. We scanned subjects as they fell asleep at night, as sleep produces major shifts in neural activity. We found that fMRI power increased during sleep at frequencies as high as 1.3 Hz, and that these power changes were correlated to EEG alpha and delta rhythms. We then developed a neural network approach to decode EEG power from fast fMRI signals, and found that alpha and delta power can be decoded from fast fMRI signals, with successful EEG prediction in subjects not included in the training set. Finally, we found that fast fMRI acquisition allowed for EEG power to be decoded from fMRI time windows as short as 0.4 s. Our results demonstrate that fMRI signals contain a wealth of high-frequency information about spontaneous neural activity that varies across arousal states.

## Results

### fMRI power as fast as 1.3 Hz increases during NREM sleep

We collected EEG and simultaneous fMRI (3T, 2.5 mm isotropic voxels, TR=378 ms) from 27 healthy adults who spontaneously transitioned between sleep and wakefulness. Decreased arousal is reflected in altered EEG rhythms: both increased EEG delta (1–4 Hz) power and decreased alpha (8–12 Hz) power (Adamantidis et al., 2019) (Fig. 1a–b). We therefore calculated EEG alpha and delta power dynamics in subjects experiencing wakefulness and progressively deeper stages of NREM sleep (N1 and N2) (Fig. 1b,c). Prior studies have shown that fMRI signals up to 0.2 Hz are correlated with neural dynamics during sleep (B. Davis et al., 2016; Fukunaga et al., 2006; Horovitz et al., 2008; Song et al., 2022). Thus, we first sought to characterize spectral fMRI changes during sleep in higher frequency ranges.

**Figure 1:**
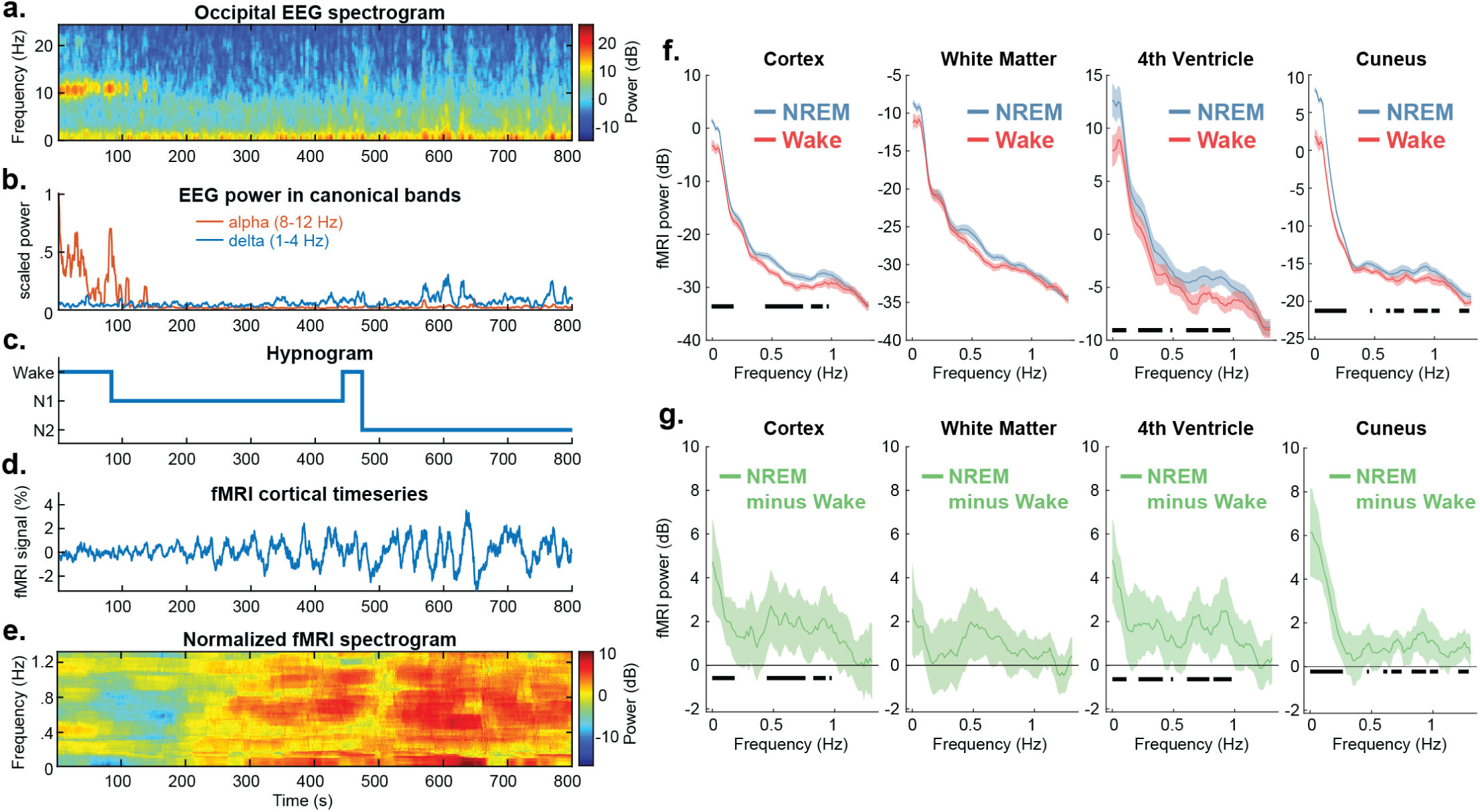
Broadband 0–1.3 Hz fMRI power increases during NREM sleep. ***a–e.*** Data from a representative subject showing a transition from wakefulness to N1 and N2. ***a.*** Occipital EEG spectrogram shows variations in coordinated neural activity across time. ***b.*** EEG power was averaged in canonical bands alpha and delta, highlighting a sharp decrease in alpha power at the start of the run, and a slow increase in delta power throughout the run. ***c.*** These EEG changes correspond to a transition from wake to N1 and N2. ***d.*** The cortical fMRI timeseries shows large-amplitude dynamics during sleep. ***e.*** Cortical fMRI spectrogram shows a broadband increase in power during sleep. The spectrogram was normalized within each frequency band to display relative changes at each frequency. ***f.*** Group-level data (*N*=23, 4 subjects excluded due to lack of stable wake segments) show significant increases in fMRI power in the cortex and fourth ventricle during NREM sleep (N1, N2, and N3) in relation to wakefulness, across most frequencies up to 1 Hz. In the cuneus (visual cortex), significant increases additionally extend to 1.3 Hz, the limit of our frequency resolution. Means with SEM. Black bars indicate significant frequencies (*p*<0.05, two-tailed paired t-tests, Benjamini-Hochberg correction). See Supplementary Fig. 1 for all brain regions. ***g.*** Group-level difference between NREM and wake fMRI power. Means of subject-wise NREM minus wake, with 95% CI. Given that baseline wake power varies substantially between subjects, we show these mean subject-wise differences to aid visualization. See Supplementary Fig. 2 for all brain regions.

We observed broadband increases in fMRI power, extending to frequencies as high as 1.3 Hz, as subjects transitioned into sleep (Fig. 1e). To quantify these spectral changes, we analyzed fMRI spectra in segments of stable wakefulness and stable NREM sleep (collapsing across N1, N2, and N3), in subjects with at least one continuous 60s segment of wakefulness (*n*=23 subjects). Due to the high sampling rate of our fast fMRI protocol (allowing detection of frequencies up to 1.3 Hz), cardiac and respiratory artifacts were not aliased, and were removed in our preprocessing pipeline. We found that fMRI power as fast as 1 Hz increased significantly in the global cortical gray matter during NREM sleep (Fig. 1f, 1g). To identify the spatial distribution of these fast fMRI effects, we anatomically parcellated individual cortical and subcortical regions (Desikan et al., 2006), and found that a significant increase in broadband high-frequency fMRI power (0.1–1 Hz) power was present in several individual cortical regions (Supplementary Fig. 1–2). The cuneus (visual cortex, Fig. 1f, 1g), displayed significant increases up to 1.3 Hz, the maximum frequency our temporal resolution could detect, suggesting it is possible that faster sampling rates could identify spectral differences in even higher frequencies. Other cortical regions also displayed significant power increases in frequencies above 1 Hz (Supplementary Fig. 1, 2). We next investigated changes in non-gray matter regions, to test whether non-neuronal fMRI signals also show an increase in high-frequency fMRI power during low arousal. We found a broadband increase in high-frequency fMRI power in the fourth ventricle (Fig. 1f, 1g), where fMRI signals have been shown to reflect fluctuations in cerebrospinal fluid (CSF) inflow (Fultz et al., 2019), indicating that high-frequency CSF inflow increases during sleep. In summary, these results demonstrated a widespread increase in high-frequency power in fMRI signals during NREM sleep, both in individual gray matter regions as well as in global signals.

### High-frequency (>0.3 Hz) fluctuations in fMRI power are correlated to EEG rhythms

Brain arousal states, and the neural activity underlying them, vary dynamically over time. We next investigated whether high-frequency BOLD signals were correlated with spontaneous neural activity, measured from fluctuations in alpha power and delta power in the EEG (Fig. 1b). These canonical neural rhythms index continuous variations in brain state: delta power increases with NREM sleep depth, and alpha power is an indicator of quiet eyes-closed wakefulness and is suppressed during NREM sleep (Adamantidis et al., 2019; Clayton et al., 2018). Subjects without resting-state alpha were excluded from alpha analyses (*N*=3, see Methods for details). We extracted alpha and delta power from occipital electrodes (which display better signal-to-noise ratio in simultaneous EEG-fMRI) in sliding 5s windows. Timepoints with subject motion or EEG artifacts were excluded prior to analysis (see Methods for details). We then calculated cross-correlations between EEG and high-frequency fMRI power (0.3–1.3 Hz, calculated in 5s windows), with a maximum lag of 40 TRs (∼15s). We did not convolve the EEG power with a hemodynamic response function (HRF), as we wanted to investigate the temporal relationships between EEG changes and high-frequency BOLD changes without assuming a specific HRF delay; however, we later repeated these analyses employing a canonical double-gamma HRF.

We found that both alpha (Fig. 2a–b, Supplementary Fig. 3) and delta (Fig. 2c–d, Supplementary Fig. 4) EEG power were significantly correlated with high-frequency (0.3–1.3 Hz) BOLD power in most of the cortex and subcortex. The positive correlations peaked at zero lag (Fig. 2) with earlier responses for higher-frequency fMRI signals (Supplementary Fig. 5), which could partially reflect shorter time delays for high-frequency hemodynamic responses. However, each rhythm also displayed unique spatiotemporal patterns of correlations reflecting rhythm-specific dynamics. BOLD power decreased around 6–10s after alpha power increased (Fig. 2a), particularly in the occipital cortex (Fig. 2b). By contrast, BOLD power increased around 0-6s after delta power increased (Fig. 2c), and these positive correlations peaked at different timings throughout the brain, with earlier timing in the frontal regions (Fig. 2d). Since fMRI signals in the fourth ventricle also varied with arousal (Fig. 1f–g), we again analyzed non-gray matter regions. EEG alpha and delta power were each significantly correlated with fMRI signals in the fourth ventricle, reflecting CSF inflow (Fultz et al., 2019), and in the white matter, in line with prior work which found BOLD responses in white matter tracts reflecting concomitant gray matter responses (Ding et al., 2025). In summary, these results establish that EEG neural oscillations are correlated with fMRI power above 0.3 Hz, representing a fast range not previously investigated in fMRI studies of vigilance states.

**Figure 2:**
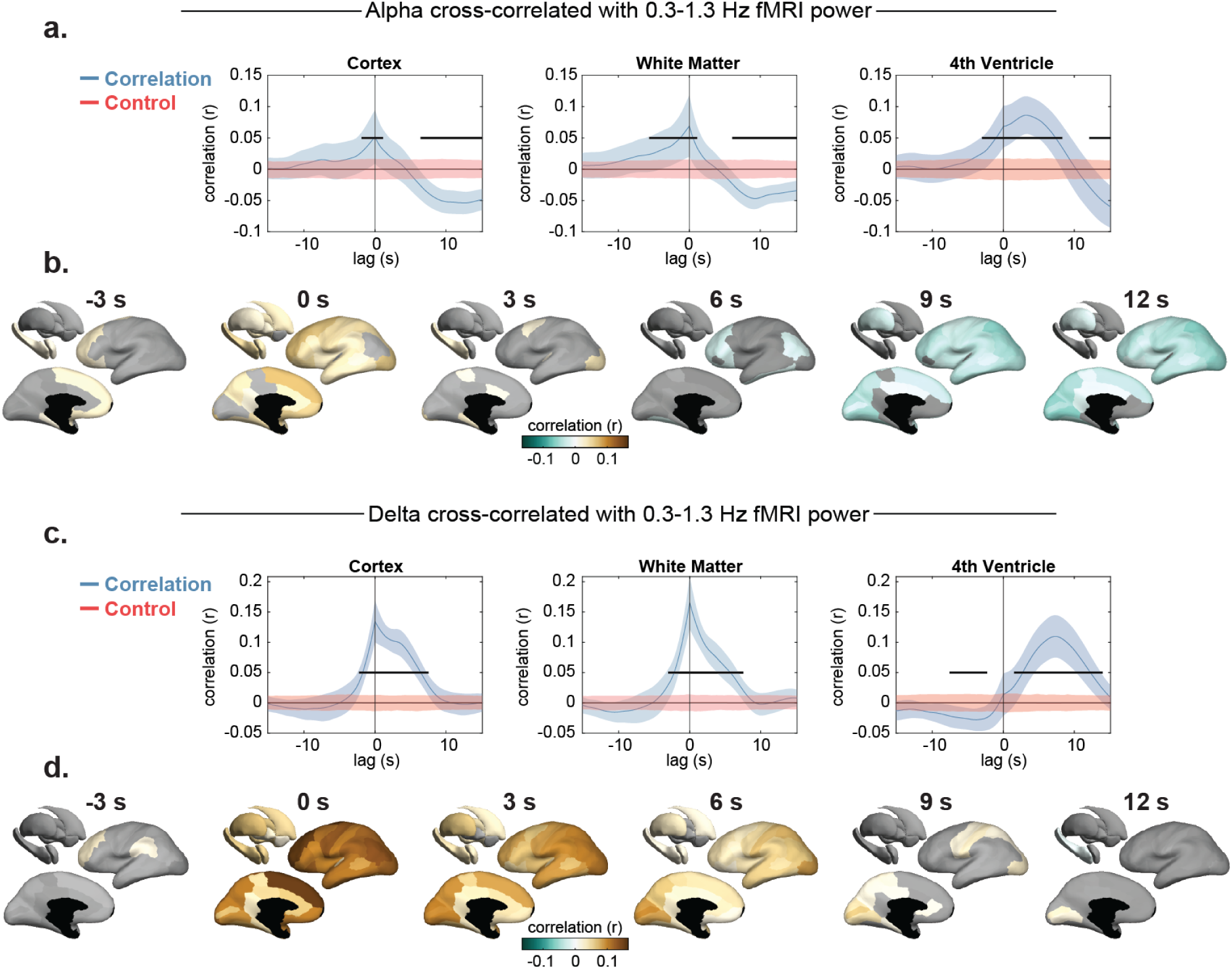
Fluctuations in high-frequency (0.3–1.3 Hz) fMRI power are significantly correlated with EEG power in the alpha and delta bands. ***a.*** Cross-correlation of high-frequency fMRI power (0.3–1.3 Hz) and EEG alpha power. Positive lag values indicate that EEG changes preceded fMRI changes. Blue lines show group means with 95% CI (*N*=24). Red lines were obtained from a randomly shifted control, and black bars indicate significant differences (see Methods for details). Full cross-correlation traces for all brain regions are shown in Supplementary Figure 3. ***b.*** Surface plots show significant correlations at specific lags between alpha power and high-frequency fMRI power in individual cortical and subcortical regions. Non-significant regions are shown in grayscale. ***c.*** Equivalent to *a.*, but for delta power (*N*=27). Full cross-correlation traces for all brain regions are shown in Supplementary Figure 4. ***d.*** Equivalent to *b.*, but for delta power.

Prior work found that narrow-band BOLD oscillations in the 0.1–0.2 Hz range are correlated with delta power (Song et al., 2022), thus we separately analyzed cross-correlations between EEG power and 0.1–0.3 Hz BOLD power (Supplementary Fig. 6–8). Cortical cross-correlation patterns were very similar to the 0.3–1.3 Hz BOLD power. Subcortical cross-correlation patterns were stronger in the 0.1–0.3 Hz BOLD range, which could be explained by the lower temporal signal-to-noise ratio in these deep brain regions, which impacts higher frequencies more strongly (Lewis et al., 2016). We also cross-correlated alpha and delta power to narrower BOLD power bands (0.1 Hz width, examining every band from 0.1 Hz to 1 Hz, including 0.1–0.2 Hz) and found that no particular band was uniquely correlated to either rhythm (Supplementary Fig. 5). In summary, the high-frequency fMRI power dynamics correlated with alpha and delta power were largely broadband, spanning the full 0.1–1.3 Hz range.

These analyses demonstrated that the power of high-frequency BOLD signals is correlated with EEG rhythms. A separate question is whether the timeseries of high-frequency BOLD oscillations is directly correlated with EEG power, which would require phase coherence at multiple frequencies between these BOLD signals and the EEG power variations. We found significant cross-correlations using the filtered BOLD signal in the 0.1–0.3 Hz range (Supplementary Fig. 9–10), but not when filtering the BOLD signal above 0.3 Hz. To ensure we did not miss any relationships that could have been identified with a traditional HRF, we conducted an additional analysis in which we convolved the EEG power trace with a double-gamma canonical HRF prior to conducting the cross-correlations (Supplementary Fig. 11). This HRF model resulted in similar correlation values between BOLD power and the EEG rhythms, and weaker correlation values between the BOLD timeseries and the EEG rhythms. These results demonstrated that both the power and the timeseries of high-frequency fMRI signals is linked to spontaneous EEG activity.

### EEG fluctuations are differentially decoded from high- and low-frequency BOLD

We next investigated whether we could decode EEG neural activity from high-frequency BOLD signals, allowing fast fMRI to be used to infer spontaneous neural activity. We hypothesized that the low signal-to-noise ratio of high-frequency BOLD, along with the unknown neurovascular coupling properties in this frequency range, would benefit from application of machine learning tools, known to operate well in low SNR regimes when functional relationships are ill-determined. A decoding approach could thus be used to reveal additional relationships between high-frequency BOLD and spontaneous neural activity.

We aimed to design a neural network capable of decoding EEG information from high-frequency fMRI. Inspired by prior work for decoding calcium imaging (Syeda et al., 2023), we used temporal convolutional layers to implicitly learn spectral information from fMRI signals (Fig. 3a). This architecture allowed the model to extract signals related to power in different high-frequency bands, while also giving it access to temporal-phase information in the BOLD timeseries. We trained the model on fMRI segments of 60 TRs (∼22 s) to predict the EEG power value at the center of the segment (Fig. 3b), choosing this mapping based on prior work (Jacob et al., 2025). We used a step size of 1 TR between windows to allow the model to decode a full continuous run of EEG alpha and delta power from the simultaneous fMRI data. In summary, the model encoded a representation of the fMRI dynamics within a 22 s window, and then translated this representation into the EEG power value at the center of the window. The decoding performance (correlation between predictions and truth) was calculated on held-out subjects that were not used to train the model nor select its hyperparameters (Fig. 3c). This prevented overfitting and ensured that the results reflect generalizable dynamics between subjects.

**Figure 3:**
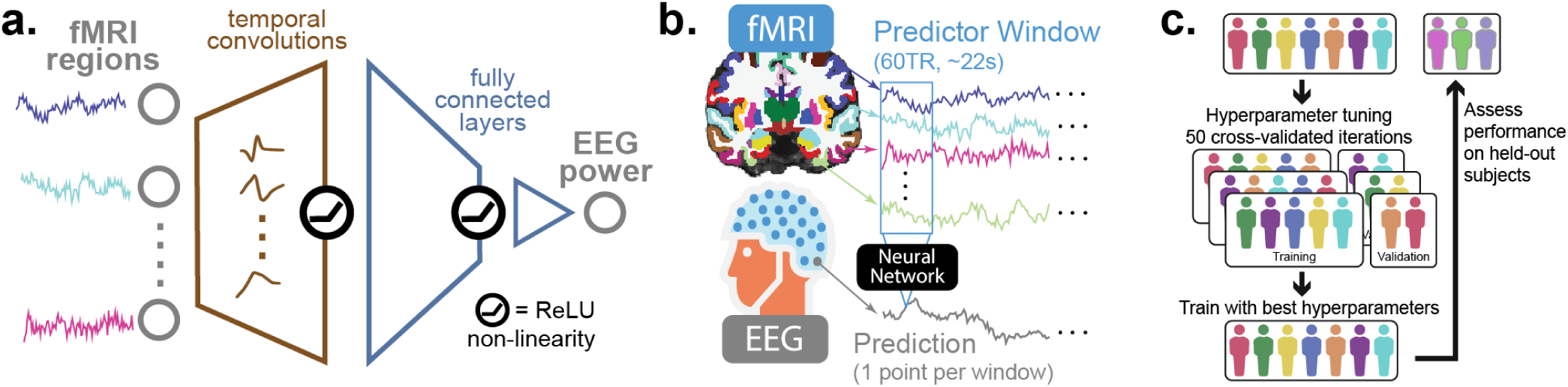
Machine learning approach for predicting alpha and delta EEG power from simultaneous fMRI data. ***a.*** We used a neural network with temporal convolutions to decode EEG power from fMRI windows. Temporal convolutional filters (learned during model training) were applied to all input fMRI regions, and the output was concatenated (across channels and timepoints) and then transformed by two fully-connected layers to predict the EEG power value. ***b.*** The model was trained on sliding windows of anatomically-parcellated fMRI data (voxels within each parcel were averaged) to predict the EEG power point (alpha or delta) at the center of the window. ***c.*** We used a train-validation-test split. Three subjects were held-out at a time, and the model used the remaining subjects to tune hyperparameters (number of convolutional filters, size of the fully-connected hidden layer, and weight decay) and train the model. Decoding performance (correlation between predictions and ground truth) was then obtained on the held-out subjects. This procedure was repeated so that a performance value was obtained for all the subjects in the dataset. We highlight that decoding performance values always reflect predictions on subject data that was not used to tune hyperparameters nor train the model.

To determine how well our approach could decode information from high-frequency components of the fMRI data, we trained models under four conditions, each using different fMRI signals as predictors (Fig. 4a): fMRI high-passed above 0.1 Hz, low-passed below 0.1 Hz, and full-spectrum fMRI, along with a control condition in which the full-spectrum fMRI data was temporally shifted randomly in relation to the EEG, breaking the relationship between the modalities (but preserving autocorrelation patterns of each modality).

**Figure 4:**
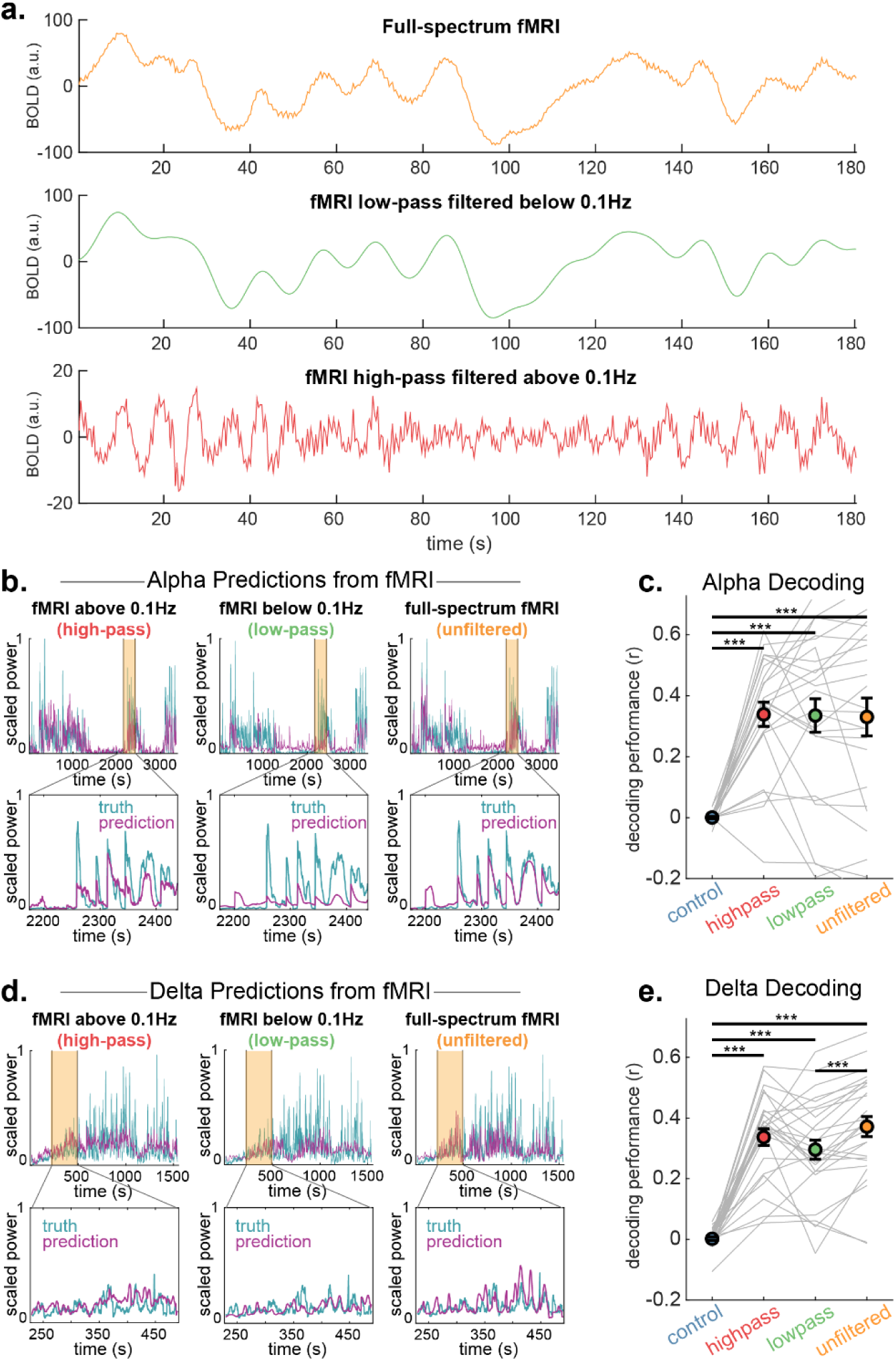
High-frequency fMRI information predicts EEG power in subjects held-out from the training set. ***a.*** We trained models using three categories of fMRI input: full-spectrum, low-pass filtered below 0.1 Hz, and high-pass filtered above 0.1 Hz. ***b.*** Example of EEG alpha power decoding on a held-out subject using the three categories of fMRI input. ***c.*** Alpha power is decoded equally well from full-spectrum fMRI, high-pass filtered fMRI, and low-pass filtered fMRI. Decoding performance was assessed via correlation between predictions on held-out subjects and ground truth. The control condition consisted of full-spectrum fMRI that was randomly shifted in relation to the EEG. *N*=24. Means with SEM. Gray lines represent held-out subjects. *** *p*<0.001, paired t-tests, Benjamini-Hochberg correction. ***d.*** Example of EEG delta power decoding on a held-out subject. ***e.*** Delta power can be decoded from fMRI above 0.1 Hz as well as from full-spectrum fMRI, but fMRI below 0.1 Hz predicts delta worse than full-spectrum. *N*=27.

We found that the network could decode high-frequency fMRI information with striking accuracy: high-frequency (>0.1 Hz) fMRI data could successfully predict EEG rhythms, and achieved comparable performance as the full-spectrum fMRI data (Fig. 4c,e). To test whether fast fMRI information contained additional information relative to conventional low-frequency fMRI signals, we compared decoding performance across filtering conditions. The low-frequency fMRI information predicted alpha power as well as full-spectrum fMRI (Fig. 4c), suggesting that high-frequency power changes carry redundant information about alpha rhythms. However, low-frequency fMRI information predicted delta worse than full-spectrum fMRI (Fig. 4e), indicating that there is unique information in fMRI signals above 0.1 Hz that predicts delta power. These results demonstrated that a neural network is capable of decoding EEG activity from high-frequency fMRI data. Importantly, the delta decoding performance results showed that high-frequency fMRI signals not only carry information about neuronal state, but also that these fast signals carry distinct information from low-frequency fMRI signals.

### EEG can be decoded from high-frequency BOLD in individual fMRI regions

To determine which brain regions contain high-frequency fMRI signatures of spontaneous neural activity, we next trained separate models on each bilateral gray matter fMRI region. In addition, to determine whether specific frequency bands are most informative about EEG alpha and delta, we high-pass filtered the fMRI signals at distinct cutoff frequencies prior to training the models.

We found that alpha-predictive information was uniformly represented across the high-frequency fMRI ranges (high-pass cutoffs from 0.3 Hz to 0.9 Hz; Fig. 5a). Delta-predictive information, on the other hand, was not decodable in fMRI frequencies above 0.9 Hz, and was weakened in the 0.6–1.3 Hz range in relation to 0.3–1.3 Hz (Fig. 5b). Most cortical and some subcortical regions could predict both alpha and delta power successfully (p<0.05, Benjamini-Hochberg corrected), likely representing global arousal dynamics, superimposed on rhythm-specific spatial variations.

**Figure 5:**
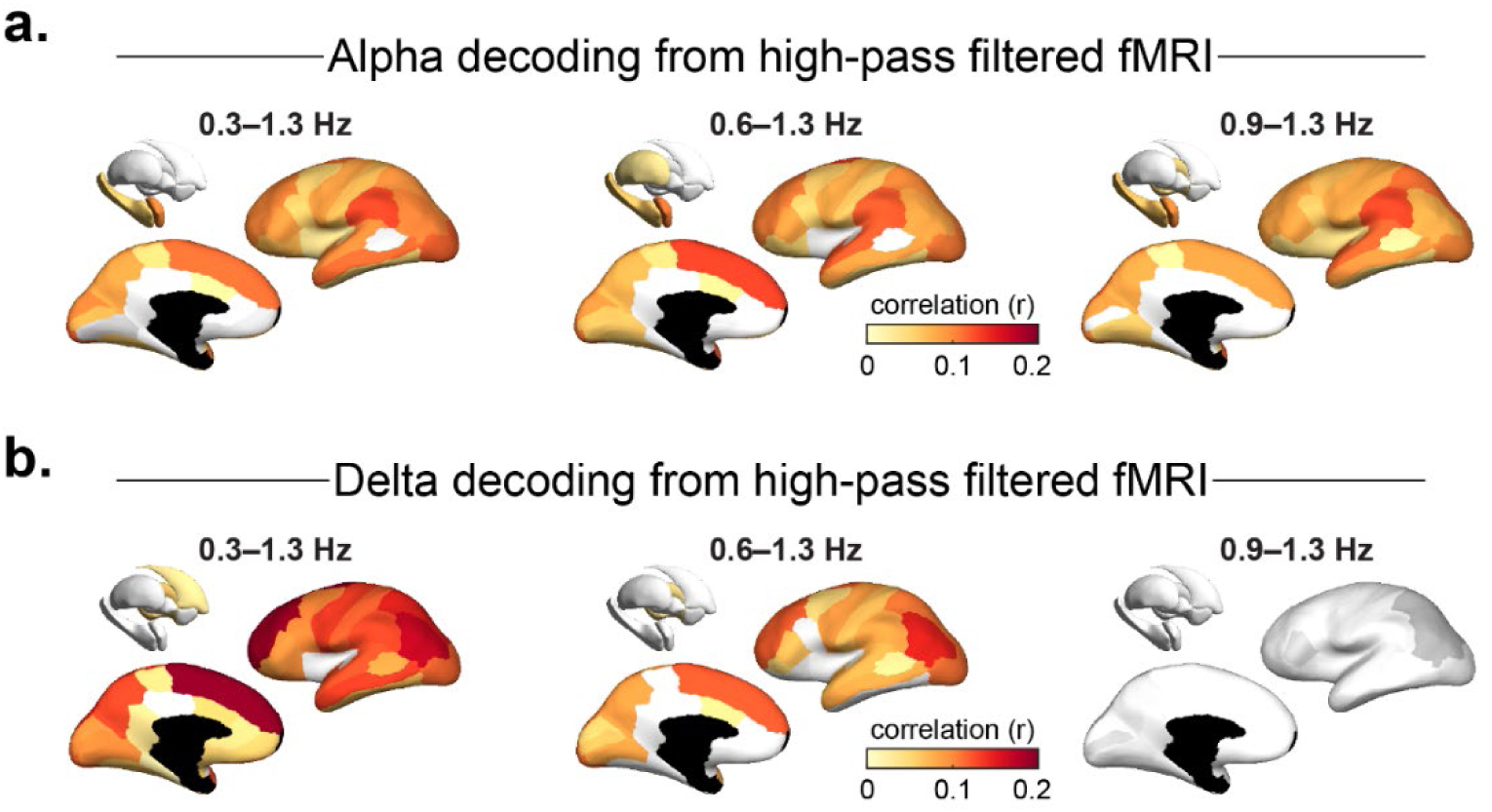
High-frequency fMRI information in each gray matter region. We trained models to predict alpha and delta power from individual fMRI regions. The fMRI timeseries of each region was high-pass filtered at different thresholds to assess at which point individual regions no longer contained high-frequency information predictive of alpha and delta. Surfaces are colored according to correlation between predictions obtained from each area and the EEG ground truth. Regions that generated non-significant predictions are shown in grayscale. ***a.*** Alpha EEG activity could be decoded from all frequency ranges investigated, from most cortical regions and some subcortical regions. *N*=24. ***b.*** Delta EEG activity was decoded most strongly from 0.3–1.3 Hz fMRI, with virtually all cortical regions providing delta-predictive information, and it could not be decoded from 0.9–1.3 Hz fMRI. *N*=27. See Supplementary Fig. 12 for region names, and performance means and SEM.

Our prior analyses showed that fMRI signals in the white matter and ventricles are significantly correlated with EEG (Fig. 1f–d, Fig. 2), demonstrating that non-neuronal fMRI signals also carry information about brain state. We therefore also assessed whether the EEG could be decoded from these signals. We found that the white matter, and to a lesser extent the ventricles, contained decodable high-frequency information, though performance was substantially worse than when decoding from the gray matter (Supplementary Fig. 13). In frequencies above 0.2 Hz, the global signal could predict delta as well as local gray matter signals (Supplementary Fig. 13b, yellow vs. blue lines), and it predicted alpha better than local signals (Supplementary Fig. 13a, yellow vs. blue lines). In summary, high-frequency fMRI signals carried information about neural rhythms both in global signals, as well as additional information in local individual gray matter regions, suggesting that both neuronal and systemic signals contribute to EEG decoding.

### Spontaneous neural activity can be decoded from sub-second windows of fast fMRI data

Since neuronal information is encoded in high-frequency fMRI signals, we hypothesized this could enable improved temporal resolution for inferring spontaneous neural activity. We therefore sought to determine whether neural activity can be decoded from very short temporal windows of fast fMRI data. Our initial decoding analyses used a 60 TR (∼22 s) window of fMRI data, aiming to predict the EEG power timepoint at the center of the window. We next trained models using progressively smaller windows of fast fMRI data, still predicting the EEG power timepoint at the center of the window. The smallest window contained a single TR of fMRI data (0.378 s). To determine whether high-frequency signals contain information beyond what is present in low-frequency signals, we trained models on full-spectrum data, low-pass filtered data (below 0.1 Hz), and high-pass filtered data (above 0.1 Hz). We additionally trained models on the global signal (using full-spectrum fMRI) to determine the impact of removing spatial information, which we hypothesized would be necessary for decoding from very short windows.

We found that alpha decoding performance was extremely well conserved across the different window lengths (Fig. 6a, yellow lines). Accurate alpha decoding from high-pass filtered fMRI data benefitted from window sizes of 10 TRs (3.8 s) or larger, consistent with the model learning high-frequency temporal dynamics from these data. Delta power suffered a decrease in decoding performance at window lengths of 7.6 s and below, but it could still be decoded well enough to obtain a mean correlation between predictions on held-out subjects and ground truth of *r=*0.28 at the 0.4–3.8 s range (Fig. 6b, yellow lines). The model leveraged information in high-frequency fMRI signals that was not present in low-frequency fMRI, evidenced by improved delta decoding from full-spectrum and high-pass fMRI compared to low-pass filtered fMRI (Fig. 6b, yellow and red versus green lines). Similar to alpha, in windows of 1.9 s and below, decoding from high-pass fMRI worsened, as there were not enough timepoints for high-frequency dynamics to be learned. Spatial information was particularly important for decoding alpha and delta from small windows, reflected in sharper drops in performance when decoding from the global signal using small windows (Fig. 6 a–b, blue lines). Prediction examples on four held-out subjects demonstrate how well alpha and delta power could be decoded from fMRI brainwide full-spectrum using 3.8 s windows (Fig. 6c–d). These results demonstrate that, when fMRI data is acquired using a fast TR, spontaneous neural activity can be reflected in the fMRI signal with enough temporal precision to allow decoding from brief time windows.

**Figure 6:**
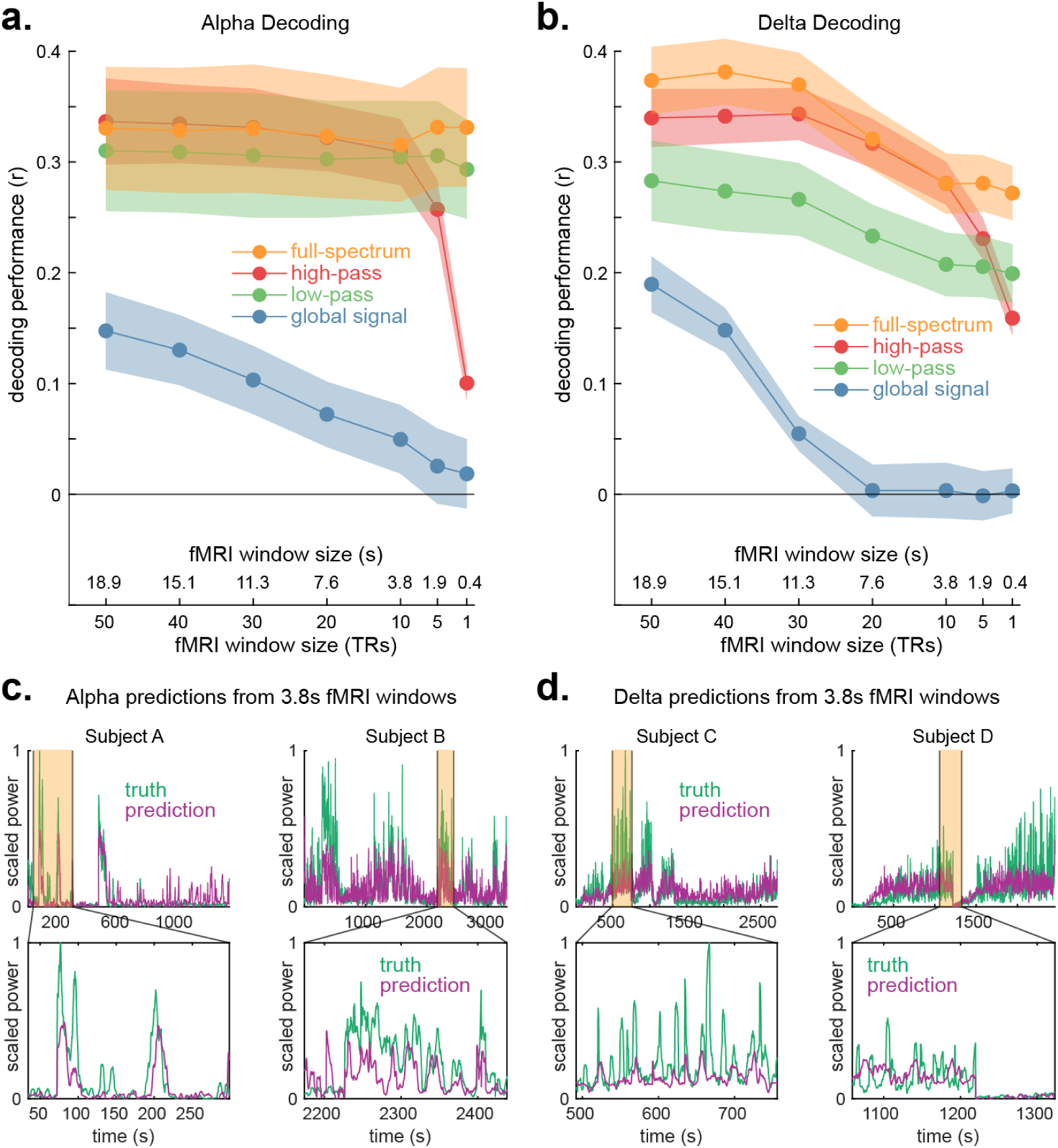
Alpha and delta power can be decoded from simultaneous fast fMRI in very short time windows, including from a single TR. ***a.*** We trained models to predict one EEG power value at the center of the fMRI window, using different window lengths and fMRI inputs: full-spectrum brainwide data, high-pass brainwide data (above 0.1 Hz), low-pass brainwide data (below 0.1 Hz), and the full-spectrum global cortical signal (removing spatial information). Decoding performance was assessed by calculating the correlation between predictions on held-out subjects and ground truth. We found that alpha power could be decoded from a single 0.378 s fMRI TR with no correlation performance deficits as long as spatial information was preserved. Accurate decoding from high-pass filtered fMRI required windows of 3.8 s or larger. *N*=24. Means with SEM. ***b.*** Delta power could be decoded from a single TR, with small performance deficits in relation to larger windows, if spatial information was preserved. Preserving high-frequency fMRI signals improved delta decoding across all window sizes, as evidenced by worse performance when using low-pass filtered fMRI in relation to full-spectrum and high-pass data. *N*=27. ***c.*** Example of alpha power decoded from short fMRI windows of brainwide full-spectrum data, on two representative subjects (who were not in the training set). The model is still able to predict long- and short-timescale variations. ***d.*** Example of delta power decoded from short fMRI windows, demonstrating the ability to predict despite small performance deficits.

## Discussion

Spontaneous neural activity profoundly shapes brain state and behavior, and measuring neural oscillations has classically required a high-temporal-resolution technique such as intracranial electrophysiology or EEG. Here, we demonstrated that fluctuations in brain states and two canonical neural rhythms tied to arousal and cognition—alpha (8–12 Hz) (Clayton et al., 2018; Pfurtscheller et al., 1996; Schneider et al., 2022) and delta (1–4 Hz) (Holz et al., 2012; Huber et al., 2004; Ngo et al., 2013; Xie et al., 2013)—were associated with clear changes in high-frequency fMRI signals. These fast fMRI components were not only significantly correlated to EEG power in the alpha and delta bands, but were robust enough to predict moment-to-moment EEG power fluctuations across subjects from subsecond windows of fMRI data. These results demonstrate that high-frequency fMRI signals carry substantial temporally precise information about spontaneous neural activity, and can be used to decode neuronal state.

### High-frequency global fMRI signals and arousal

EEG alpha and delta both strongly fluctuate with arousal, and reflecting this commonality, both rhythms were correlated with, and could be decoded from, high-frequency BOLD activity in most cortical regions, along with the global cortical signal. Thus, the coupling of global fMRI signals and EEG rhythms was found to be a substantial contributor to how high-frequency fMRI patterns represent spontaneous neural activity. Recent work has demonstrated that seconds-scale metrics of global arousal can explain much of the variance present in slow neural activity (Raut et al., 2025). Slow variations in global BOLD dynamics, on the order of tens of seconds, are also known to reflect cyclic shifts in arousal (Bolt et al., 2025; Falahpour et al., 2018; Goodale et al., 2021; Liu et al., 2018; Raut et al., 2021; Zhang et al., 2023). Here, we demonstrated that global arousal variations modulate not only conventional low-frequency BOLD signals, but also affect high-frequency BOLD contents, with tighter timing relationships in the order of hundreds of milliseconds.

### Distinct high-frequency BOLD signatures of alpha and delta

Beyond global arousal dynamics, high-frequency fMRI signals also contained region- and frequency-specific information about alpha and delta. These spatiotemporally distinct relationships could reflect the circuits specifically engaged in these neural rhythms. Alpha-specific signatures included significant anti-correlations with high-frequency BOLD power that were stronger in the occipital cortex (Fig. 2b). Alpha rhythms covary with changes in visual processing (see Clayton et al. (2018) for a review), and prior EEG-fMRI work found that BOLD signal in the occipital cortex is anticorrelated with alpha power (de Munck et al., 2007; Feige et al., 2005; Goldman et al., 2002; Martínez-Montes et al., 2004; Moosmann et al., 2003; Tyvaert et al., 2008). Our results show that alpha-coupled visual cortex deactivation may also induce a decrease in high-frequency BOLD responses.

Delta power changes can represent multiple sleep oscillatory dynamics, including slow (0.1-1 Hz) oscillations and slow (0.5-4 Hz) waves, capturing their collective frequency and amplitude. The frequency range that our fMRI sampling rate could resolve (<1.3 Hz) overlaps with the range of these oscillatory dynamics, and thus high-frequency BOLD signals could be particularly valuable by providing a temporally precise signature of delta dynamics. Supporting this hypothesis, high-frequency BOLD activity (>0.1 Hz) contained information about delta that was not present in low-frequency BOLD components (<0.1 Hz) (Fig. 4e and Fig. 6b, yellow vs. green points). Delta power was also linearly correlated with high-frequency BOLD power (Fig. 2c), and the cross-correlation patterns were spatiotemporally structured, with stronger early positive correlations in the anterior portion of the brain, and a shift towards the occipital cortex at later timepoints. This pattern is aligned with electrophysiological work that showed oscillations in the delta range are most commonly initiated in the frontal part of the brain, and travel towards posterior regions (Betta et al., 2021; Massimini et al., 2004), again suggesting that high-frequency BOLD signals can contain precise signatures of delta dynamics.

### High-frequency fMRI signals outside of the gray matter

Signals outside of the gray matter are often seen as irrelevant systemic signals, or ‘physiological noise’. However, recent work has shown that the white matter displays BOLD signals consistent with its fiber tract architecture and concomitant gray matter responses (Ding et al., 2018). Additionally, fMRI signals caused by cerebrospinal fluid flow in the ventricles, which can be detected with fast fMRI acquisition paradigms as used here, can be modulated by synchronous neural activity (Fultz et al., 2019; Williams et al., 2023). Finally, similar to global arousal, systemic physiology can carry important information about neural activity (Bolt et al., 2025; Bright & Murphy, 2015; Chen et al., 2020; Driver et al., 2016; Kandimalla et al., 2025). In this study, we found that high-frequency fMRI signals in the global white matter displayed cross-correlation patterns with EEG power that closely resembled the patterns of the global cortical signal. This observation is in line with prior studies, which found that white matter BOLD signals show similar information as the gray matter, including information about stimuli (Mishra et al., 2020), reaction times (Yarkoni et al., 2009), and spontaneous neural activity (Ding et al., 2025; Huang et al., 2023); our work demonstrates that high-frequency gray matter dynamics are also reflected in the white matter. Conversely, the white matter displayed smaller power differences between sleep and wakefulness than the cortex. Therefore, global cortical fMRI signals are likely reflected in the white matter with similar temporal patterns, but lower amplitudes.

We also found significant relationships between fMRI signals in the fourth ventricle and the EEG rhythms, demonstrating that this fluid physiology carries information about neural activity. Global cerebral blood volume changes can drive cerebrospinal fluid (CSF) flow in the fourth ventricle, by mechanically displacing CSF (Fultz et al., 2019; Williams et al., 2023). Furthermore, EEG power is known to be correlated with low-frequency (<0.1 Hz) CSF signals (Yang et al., 2025). We found positive correlations between fMRI power above 0.3 Hz in the fourth ventricle with alpha and delta power, and the peak of these correlations was later than the peak of the correlations between the global cortical signal and the EEG. This coupling is consistent with cerebral blood volume changes driving CSF flow, and demonstrates that this dynamic can also happen in frequency ranges above 0.1 Hz, with CSF rapidly being displaced during neural and vascular dynamics.

### Temporally precise decoding of neural activity

Fast fMRI sampling has the potential to enable the investigation of neural dynamics with greater temporal precision. We found that we could decode spontaneous neural activity from a single fMRI TR of 378 ms aligned with the center of the predicted EEG power value (acquired in a 5 s window). Prior work has shown that stimulus-induced neural activity can be decoded from shorter windows earlier than the peak of the hemodynamic response (Kohler et al., 2013) using a TR of 739ms and a linear decoder, with peak decoding achieved as fast as 3 seconds after stimulus onset, and above chance decoding achieved after 2 seconds. More recent work acquired 7T fMRI data at a very fast sampling rate (TR=125ms), and achieved above-chance decoding from a 2 second window post-stimulus, also using a linear decoder (Miyawaki et al., 2025). Interestingly, this prior work used temporally smoothed fMRI data, which improves SNR but removes high-frequency components. Given our finding that low-pass filtering fast fMRI signals can result in loss of decodable information (see Fig. 4e and Fig. 6b), and that high-frequency fMRI information contains non-linear components, future work could investigate whether stimulus decoding may be possible with even greater temporal precision if high-frequency information is preserved and a non-linear model is used.

### Potential mechanisms of high-frequency hemodynamic responses to neural activity

The presence of high-frequency hemodynamic information representing spontaneous neural activity raises questions about the underlying mechanisms of neurovascular coupling leading to these signals. One possibility is that fast, non-linear hemodynamic responses (Polimeni & Lewis, 2021) underlie these high-frequency signals. While our imaging protocol was not designed to specifically isolate responses from the microvasculature, which are thought to be particularly temporally precise (Gomez et al., 2024; Siero et al., 2011; Uludağ & Blinder, 2018), these are nonetheless present in our data. A portion of these signals would be lost due to their lower amplitude and temporal phase cancellation effects, but a neural network model may be capable of recovering enough information about high-frequency hemodynamic responses in the microvasculature to achieve the observed EEG decoding. High-frequency BOLD power was also linearly correlated to spontaneous neural activity. A portion of this relationship is likely explained by global variations in arousal, which could modulate the neural rhythms as well as the vasculature. However, given that correlation patterns were also spatially structured throughout the brain differently for each rhythm (Fig. 2), they likely also represent specific, local responses related to alpha and delta. Ultimately, future work is necessary to determine the vascular mechanisms underlying these high-frequency BOLD responses.

### Limitations and future perspectives

We revealed distinct spatially structured patterns of high-frequency fMRI information underlying each EEG rhythm, though the frequency range was limited by our sampling rate and by our 3T acquisition, since high-frequency BOLD signals are small and difficult to detect. Ultra-high field fMRI acquisition and faster sampling rates could allow for additional information to be identified. For instance, we found that the cuneus (Fig. 1f, 1g) displayed significant power increases during sleep in relation to wake up to 1.3 Hz, which was the limit that could be resolved by our sampling rate; these significant changes might thus extend to faster frequency ranges. The improved signal-to-noise ratio provided by ultra-high field fMRI (Setsompop et al., 2016) could also allow for higher-frequency information to be identified in other regions, particularly deep brain regions. Finally, we show that combining fast fMRI with a neural network decoding approach enables information about neural activity to be inferred from very short fMRI time windows. This can allow for more naturalistic experimental designs, shorter inter-stimulus intervals (and thus a greater number of trials) during acquisition, and a greater ability to separate distinct neural signatures in fMRI data.

### Conclusions

We found that spontaneous neural activity measured with EEG is coupled to high-frequency fMRI activity, reflecting both global arousal dynamics, and fMRI patterns unique to distinct neural rhythms. This high-frequency fMRI activity was robust enough to allow for machine learning-based EEG decoding in subjects that were not present in the training set. Our decoding approach demonstrated that high-frequency BOLD signals contains unique neurally-coupled dynamics not present in the slower BOLD range. Many fMRI analysis pipelines involve low-pass filtering the fMRI signal, aiming to remove noise. While noise is certainly present, our results reveal that this common processing step may also be removing important information about neural activity. In summary, high-frequency fMRI signals contained rich and temporally-precise signatures of spontaneous neural activity.

## Methods

### Participants

Data analyzed in this work was collected as part of a different study (Bailes et al., 2025). Only resting-state data from the cohort of younger subjects (ages 18–40) who completed a nighttime MRI session were used in this work.

Data acquisition was performed at two sites. All subjects provided written informed consent. At Site 1, procedures were approved by the Boston University Charles River Campus Institutional Review Board (IRB #6031). At Site 2, procedures were approved by the Massachusetts Institute of Technology Institutional Review Board (IRB #23030000951). Exclusion criteria included the regular use of sleep aid medications, current psychiatric or neurological disorders, cognitive impairment, any major medical disorders, sleep disorders, substance use disorders, excessive (estimated >=400 mg daily) caffeine intake, or any MRI contraindications.

Twenty-eight subjects completed an MRI session (Site 1: *n*=20; Site 2: *n*=8). Night scans were scheduled to begin around midnight, with the exact start time depending on factors related to the equipment set up and preparation of EEG caps. Subjects were instructed not to consume any caffeine after 11:00 AM on the day of the experiment and not to nap the day of the experiment. Each subject was scanned for 1–3 runs of sleep opportunity, with each run lasting 25 minutes. The number of runs depended on factors such as experimental setup time and subject comfort. One subject was excluded due to high motion in all runs. Twenty-seven subjects were analyzed in this study (mean age 23.7, 18–39 range, 15 female, 12 male).

### EEG-fMRI acquisition

At both sites, subjects were scanned on a 3T Siemens Prisma scanner with a 64-channel head and neck coil. Pupil tracking (used here to confirm that eyes were closed) was conducted with an EyeLink 1000 Plus Eyetracker (SR Research). Anatomical images were acquired using a 1 mm isotropic T1-weighted multi-echo MPRAGE (van der Kouwe et al., 2008). Functional runs were acquired using a Simultaneous Multi-Slice (Setsompop et al., 2012) single-shot 2D gradient-echo EPI sequence (TR=0.378 s, TE=31 ms, 2.5 mm isotropic voxels, 40 slices, Multiband factor=8, blipped CAIPI shift=FOV/4, flip angle=37°, no in-plane acceleration). The bottom functional edge slice was placed so that it intersected the fourth ventricle, which allowed for the acquisition of cerebrospinal fluid inflow signals (Fultz et al., 2019). Pulse oximetry, skin conductance, and respiration were monitored with BIOPAC sensors (BIOPAC Systems).

At both sites, EEG was acquired using MR-compatible 32-channel EEG caps fitted with 4 carbon-wire loops (BrainProducts GmbH, Germany) at a sampling rate of 5000 Hz. EEG acquisition was synchronized to the MRI scanner 10 MHz clock to reduce aliasing of high-frequency gradient artifacts. Additional sensors were used to record systemic physiology: respiration was measured concurrently using an MRI-safe pneumatic respiration transducer belt around the abdomen and pulse was measured with a photoplethysmogram (PPG) transducer (BIOPAC Systems, Inc., Goleta, CA, USA). Physiological signals were acquired at 2000 Hz using Acqknowledge software and were aligned with MRI data using triggers sent by the MRI scanner. Two built-in electrooculography (EOG) electrodes were used to record eye movements. Signals from three electromyography (EMG) electrodes were recorded and one was selected for sleep stage scoring.

### fMRI pre-processing and parcellation

Functional runs were slice-time corrected using FSL version 6 (slicetimer; https://fsl.fmrib.ox.ac.uk/fsl/fslwiki), (Jenkinson et al., 2012) and motion corrected to the middle frame using AFNI (3dvolreg; https://afni.nimh.nih.gov/). Physiological noise was removed using HRAN, open-source software that implements a statistical model of harmonic regression with autoregressive noise (Agrawal et al., 2020) (https://github.com/LewisNeuro/HRAN). Motion corrected data were registered to each individual’s anatomical data using boundary-based registration as implemented in FreeSurfer (bbregister, https://surfer.nmr.mgh.harvard.edu/) (Greve & Fischl, 2009).

We highlight that our physiological noise correction removed noise associated with specific cardiac and respiratory cycles, but did not regress out the low-frequency component of cardiac and respiratory rate variations, as regressing these low-frequency signals can also remove information about neural activity that is coupled to systemic physiology (Driver et al., 2016; Kandimalla et al., 2025).

fMRI parcellation was performed in FreeSurfer based on the Desikan-Killiany atlas (Desikan et al., 2006). The following region labels were obtained: 31 bilateral cortical regions (the entire cortex minus the temporal pole, frontal pole, and entorhinal regions, which were not covered by the acquisition volume in all subjects), 7 bilateral subcortical regions (thalamus, pallidum, amygdala, caudate, putamen, accumbens, hippocampus), and 8 non-gray matter regions (left- and right- total white matter, left- and right- lateral and inferior lateral ventricles, third ventricle, and fourth ventricle); total of 84 regions. Anatomical labels were interpolated into functional space, and functional voxels with 70% or above of overlap with the label were averaged to generate the mean region time series. The first 20 TRs of each run were discarded to remove transient artifacts as the MRI signal reached steady-state.

### EEG pre-processing

Gradient artifacts were removed using average artifact subtraction (Allen et al., 2000) with a moving average of the previous 20 TRs. Ballistocardiogram artifacts were removed from each EEG channel using signals from the 4 carbon-wire loops using the sliding Hanning window regression method from the EEGLab CWL toolbox (van der Meer et al., 2016) with the following parameters: window=25 s and lag=0.09 ms. Cleaned EEG signals were re-referenced to the average of EEG channels.

### Sleep scoring

Two trained scorers manually sleep scored EEG data in 30-second epochs according to the AASM sleep scoring guidelines (Berry et al., 2017) using the Visbrain GUI (https://pypi.org/project/visbrain/) and EEG and EOG channels that were bandpass filtered between 0.3–20 Hz. Any discrepancies between the two scores were discussed and merged before final analysis. A breakdown of time spent in each sleep stage across all subjects is shown in Supplementary Fig. 14.

### State-based spectral power analysis

For identifying spectral differences between wake and sleep, the power of fMRI signals was calculated using multi-taper spectral estimation in Python using the spectral_connectivity toolbox (Denovellis et al., 2022; Thomson, 1982). Power was estimated in 60-second segments of continuous wakefulness or NREM sleep, and any 60-second segments that had average framewise displacement > 0.3 mm or maximal framewise displacement above 0.75 mm were excluded from analyses. Pairwise comparisons in each spectral frequency across sleep and wakefulness were performed in subjects that had both wake and NREM sleep data (*N*=23) using a paired t-test, and were corrected for multiple comparisons using the Benjamini-Hochberg procedure.

### Coherence analysis

We used the *coherencyc* function from the Chronux toolbox (Bokil et al., 2010) in MATLAB (Mathworks, Natick, MA) to calculate the multi-taper coherence between pairs of BOLD timeseries. Coherence was calculated between 60-second non-overlapping motion-free epochs. A framewise displacement exclusion threshold of 0.2 mm was used to exclude epochs with motion.

### Cross-correlation analyses

#### EEG-fMRI power acquisition

EEG and fMRI power were calculated via the multi-taper spectrogram method (Thomson, 1982) using an open-source implementation in MATLAB (Prerau et al., 2017). Spectrograms for each modality were obtained using a 5 second window, centered on each fMRI TR. For the Supplementary analyses where we examined BOLD power in the 0.1–0.3 Hz range, spectrograms were instead obtained using 20s windows, as a 5s window would be unable to resolve this slower range. A 20s window was also used for the analysis in Supplementary Fig. 8.

For EEG power measurements, we used occipital electrodes as they contain less MR-induced noise and improved data quality due to being pressed against the back of subjects’ heads as they lie down in the scanner. EEG spectrograms were visually inspected for the presence of clear eyes-closed alpha rhythms. Most healthy adults display a clear increase in alpha power when awake with eyes closed (see Fig. 1a), but a small percentage of individuals do not show this increase for reasons not yet understood (Barzegaran et al., 2017; H. Davis, 1936; H. Davis et al., 1938; Niedermeyer, 1999). As this difference may reflect distinct underlying neural dynamics, we excluded these subjects from the alpha analyses (*N*=3 subjects were excluded). These subjects were included in the delta analyses. The EEG power time series was then extracted for the frequencies of interest: 1.1–4 Hz for delta (chosen to minimize contamination from low-frequency artifacts in EEG-fMRI) and 8.5–12 Hz for alpha (chosen to minimize contamination from theta and spindle frequency ranges while still covering subject variations in alpha range). Power in the 40–52 Hz range was also extracted for the purpose of artifact detection (discussed below). Power was extracted from three occipital electrodes (O1, O2, and Oz) and then averaged. For the analyses involving HRF convolution, we convolved the EEG power with a double gamma HRF with a peak time equal to 6 seconds.

#### Data exclusion criteria

To mark data points unsuitable for analyses, the following criteria were used: 1) TRs with motion above 0.3 mm; 2) z-scored EEG power in the 40–52 Hz range with a value above 1 (adjusted to be lower for runs with particularly noisy data; determined qualitatively for each subject prior to all analyses, via visual examinations of EEG spectrograms); 3) detected EEG local outliers (distance larger than 6 local scaled median absolute deviation (MAD)) within a sliding window of 1000 TRs (thresholds were reduced to 3 MAD for subjects with excessively noisy data). The outlier detection method was applied to delta power and power in the 40–52 Hz range (but not to alpha power, as its naturally fast fluctuations would be erroneously detected as outliers). Datapoints that fulfilled any of these criteria were then removed, and the remaining data were saved as separate segments to avoid creation of spurious continuities.

#### Cross-correlation and statistical testing

Within each subject, cross-correlations were calculated on each segment, with a maximum lag of 40 TRs. A weighted average, using segment length as the weight, was calculated to obtain the results for each subject. As a control, the EEG time series was shifted in relation to the fMRI by a random value for each subject prior to obtaining the cross-correlations, and this procedure was repeated 1000 times to generate a null distribution. A p-value was then assigned to each lag based on which percentile of this null distribution equated to the group mean of the cross-correlation lag (e.g., if the mean was equivalent to the top 1%, this would be equal to a p-value of 0.02 for a two-tailed procedure). The p-values were then corrected for multiple comparisons using the Bonferroni procedure.

### Machine learning methods

#### Data preparation

The averaged fMRI timeseries within each parcel (separately for each subject) was z-scored within each run. EEG power for each subject (combining runs, if multiple runs were acquired for a given subject) was scaled to 0–1, separately for alpha and delta. For analyses that involved filtering the fMRI data, Butterworth filters were applied prior to z-scoring and segmentation.

fMRI data were split into overlapping 60-TR sections, with the goal of predicting the EEG power at the point in the center of the fMRI window (equivalent to the position of the 30^th^ fMRI point). We predicted the EEG power at this point at the center of the window to avoid assumptions about temporal relationship between the two modalities (such as regions which might have early or late responses).

In order to predict every EEG point that had been interpolated to match the fMRI temporal sampling grid, there was a 59-TR overlap between each adjacent predictor window (windows were slid by 1 TR). Given that no fMRI samples would be available for EEG points at the edges of the data segments following artifact removal, the first 29 and the last 30 EEG points of each segment were discarded.

#### Neural network methods

Our neural network was implemented in Pytorch (https://pytorch.org/). It consisted of a temporal convolutional layer (Conv1d; number of filters tuned between 8 and 64, and kernel of size 21) followed by a ReLU non-linearity, a fully-connected layer (size tuned between 8 and 64), another ReLU non-linearity, and a final fully-connected layer that generated a single scalar output value (the EEG power at the center of the fMRI window). The convolutional kernels are learned from the data along with the remaining model weights, allowing the relationship between the fMRI and EEG to be determined in a data-driven way. L_2_ regularization was tuned between 0.001 and 0.01 (these values were selected after early testing of the model showed that it benefitted from strong regularization). A batch size of 32 was used, and at the end of each epoch the learning rate (Adam learner, starting rate of 0.001 (Kingma & Ba, 2017)) was divided by a factor of 10 (learning-rate annealing).

We allowed the model to fit a subject-specific bias value (a fixed number added to all predicted points for that specific subject). This value allowed for the full decoded time course to be ‘shifted’ up and down, which did not affect our decoding performance metric (Pearson’s correlation), but improved model learning (as the model was trained to minimize the mean squared error between predictions and ground truth). This subject-specific bias can be interpreted as accounting for variations in mean EEG amplitudes, which are partly derived from the specific arousal states that a subject exhibited during data acquisition. For the held-out subjects, this bias was fit using 5-fold cross-validation.

#### Model training and evaluation

We used a train-validation-test split. At each training and tuning fold, all data for three subjects were held out as test data. The remaining subjects were used for 50 iterations of hyperparameter tuning. At each iteration, 5-fold cross validation was applied so that the model using that iteration’s specific set of hyperparameter values could be evaluated on all training subjects. The tuned hyperparameters and their ranges are described above. Tuning procedures used early stopping, ceasing training if an epoch’s performance was worse than the prior epoch. Early stopping was also used to determine the optimal number of epochs, with 5 as the maximum (a low number was used due to our sliding-window approach which used a step size of 1; in other words, the fMRI data between neighboring windows was very similar, so a larger number of epochs would have resulted in overfitting). After 50 iterations, the best hyperparameters were selected, and all training subjects were used to train the model with the optimal number of epochs. The trained model was then used to generate predictions on the held-out subjects, and decoding performance (correlation between predictions and truth) was calculated. In other words, the held-out subjects were never used either for hyperparameter tuning or model training.

The tuning procedure was skipped due to computational resource constraints for the analyses shown in Fig. 5c–d and in Fig. 6. For these analyses, the number of convolutional filters was set to 32, the number of epochs was set to 4, the L2 regularization was set to 0.001, and the fully-connected layer size was set to a fifth of the number of brain regions provided to the model (rounded down), with 8 as the minimum size (in practice, this equated to a size of 8 for Fig. 5c–d, and 16 for Fig. 6). These numbers were selected as they were the original hyperparameters used prior to the implementation of tuning; they were selected through manual exploration on a separate dataset.

We highlight that normalization (see ‘Data preparation’ above) was performed separately within each subject so that at no point did the held-out subject data affect the training data. After training the model, its weights were fixed and the Pearson’s linear correlation coefficient was calculated between the predictions made for the held-out subjects and the ground truth.

In order to achieve more precise estimates of decoding performance, we repeated each fold 10 times with distinct random seeds, yielding 10 trained models and thus 10 sets of predictions (plus 10 Pearson’s correlation values) for each held-out subject. In all analyses, each correlation value reported for each held-out subject represents the mean correlation performance of 10 models.

## Acknowledgements

This study was funded by National Institutes of Health grants R01-AG070135, R01-EB019437, U19-NS128613, and U19-NS123717; the Sloan Fellowship, the McKnight Scholar Award, the Pew Biomedical Scholar Award, the Simons Collaboration on Plasticity in the Aging Brain (811231). Resources were provided by NSF Major Research Instrumentation grant BCS-1625552. Collaboration was facilitated by the Janelia Visiting Scientist Program.

