## Supplementary figures for "Functional MRI signals at and beyond 1 Hz are coupled to brain states and predict spontaneous neural activity"

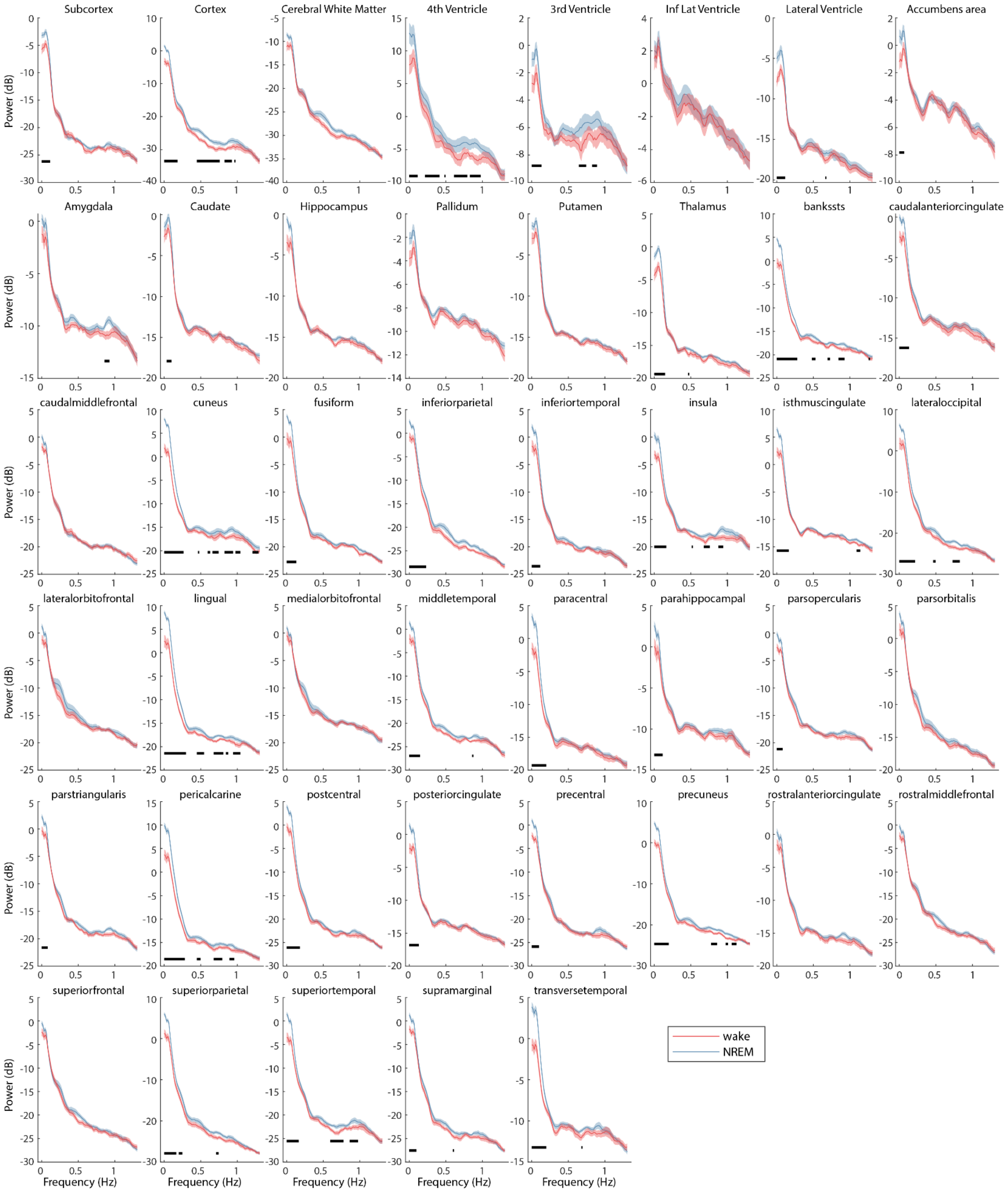


**Supplementary Figure 1:** fMRI power increases in NREM sleep in relation to wake in several brain regions. Means with SEM, N=23. Black bars indicate significant frequencies (p<0.05, two-tailed paired t-tests, Benjamini-Hochberg correction).

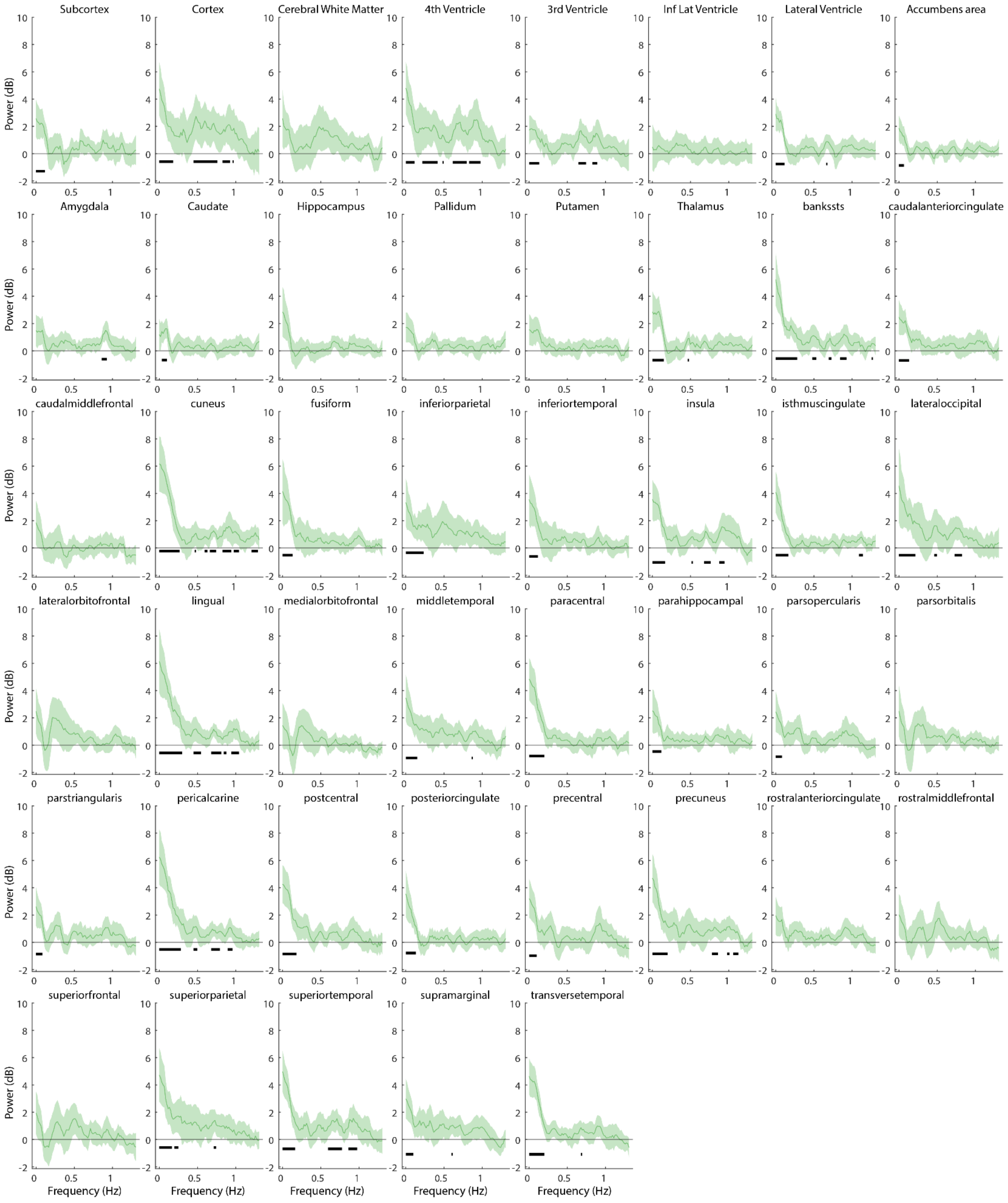


**Supplementary Figure 2:** Group-level (N=23) NREM minus wake fMRI power for each brain region; given that baseline wake power varies substantially between subjects and brain regions, we present the mean subject-wise differences to aid visualization (means with 95% CI). Black bars indicate significant frequencies ($p<0.05$, two-tailed paired t-tests, Benjamini-Hochberg correction).

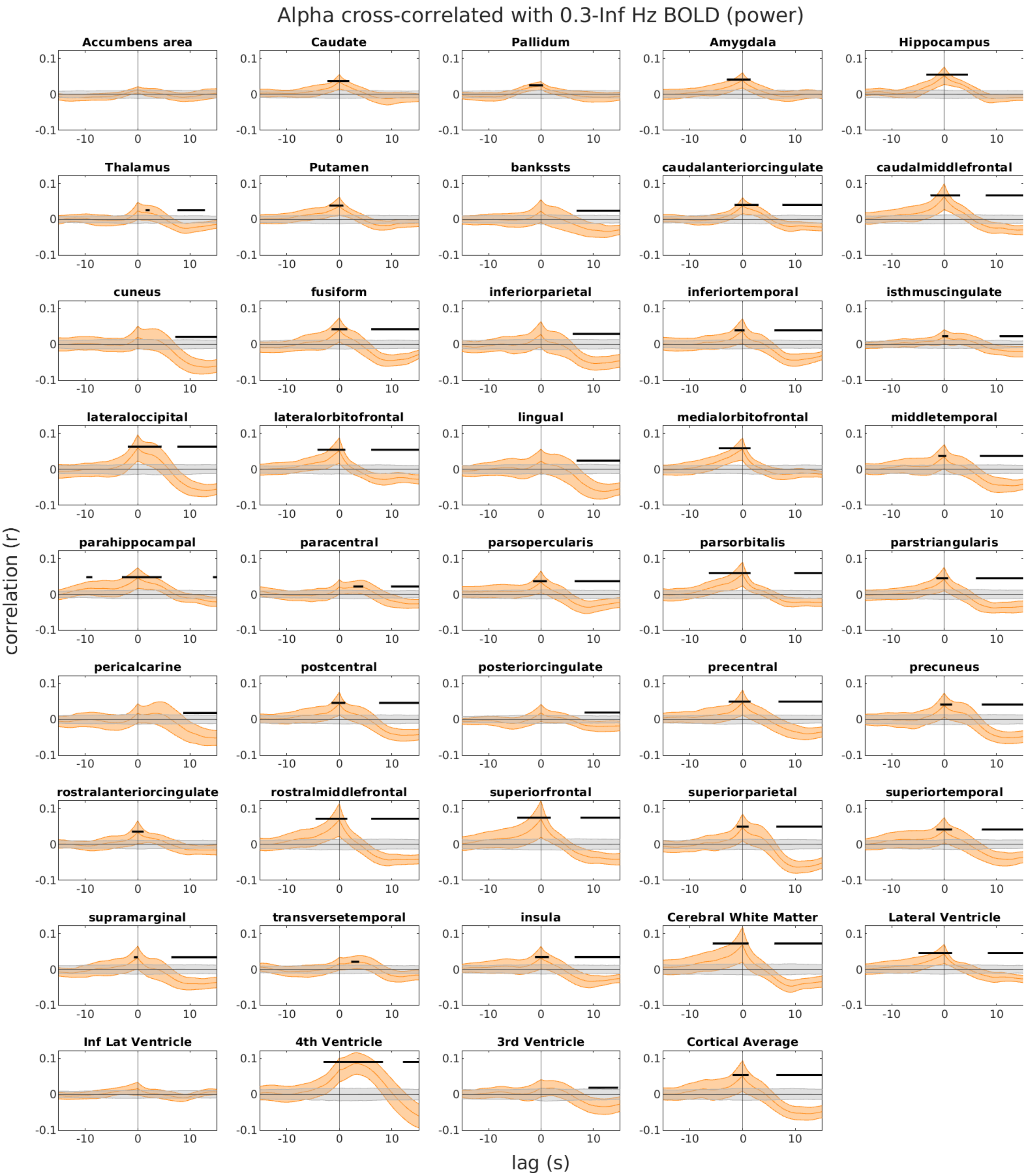


**Supplementary Figure 3:** Alpha power is significantly correlated with fMRI power above 0.3Hz in virtually all regions. Patterns differ throughout the brain; see Figure 2b for cortical surface plots. Generally, a positive correlation around lag zero (where the power windows for both EEG and fMRI were centered) was followed by a negative correlation at a positive lag (indicating that fMRI power decreased around 6-10s after increases in EEG alpha power). Yellow lines show group means with 95% CI. Gray lines were obtained from a randomly shifted control, and black bars indicate significant differences (see Methods for details). N=24.

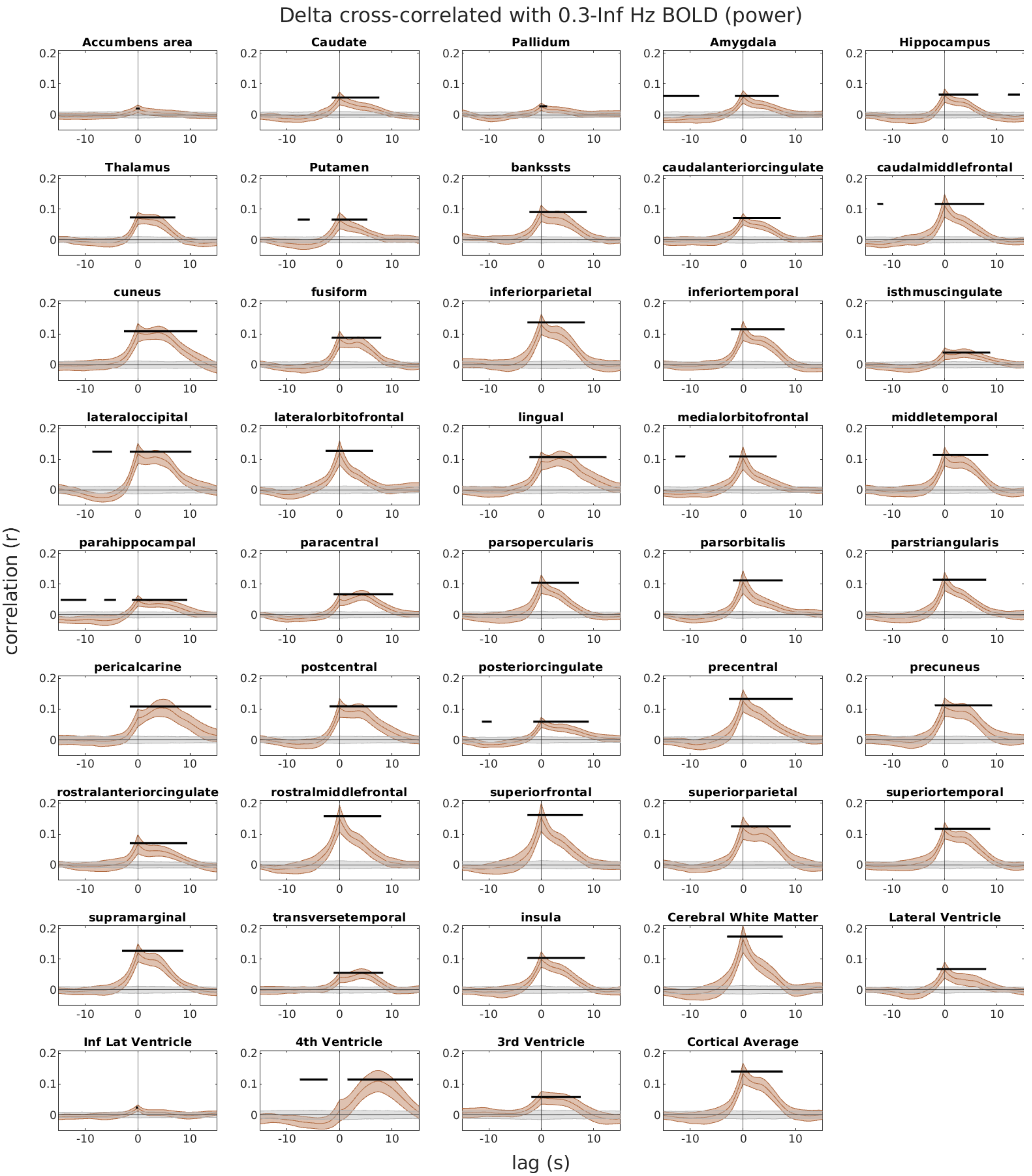


**Supplementary Figure 4:** Delta power is significantly correlated with fMRI power above 0.3Hz in all regions. Patterns differ throughout the brain; see Figure 2d for cortical surface plots. Correlations between fast fMRI power and delta were overall stronger than with alpha, and generally only positively correlated. Positive lag values indicate that EEG changes preceded fMRI changes. Brown lines show group means with 95% CI. Gray lines were obtained from a randomly shifted control, and black bars indicate significant differences (see Methods for details). N=27.

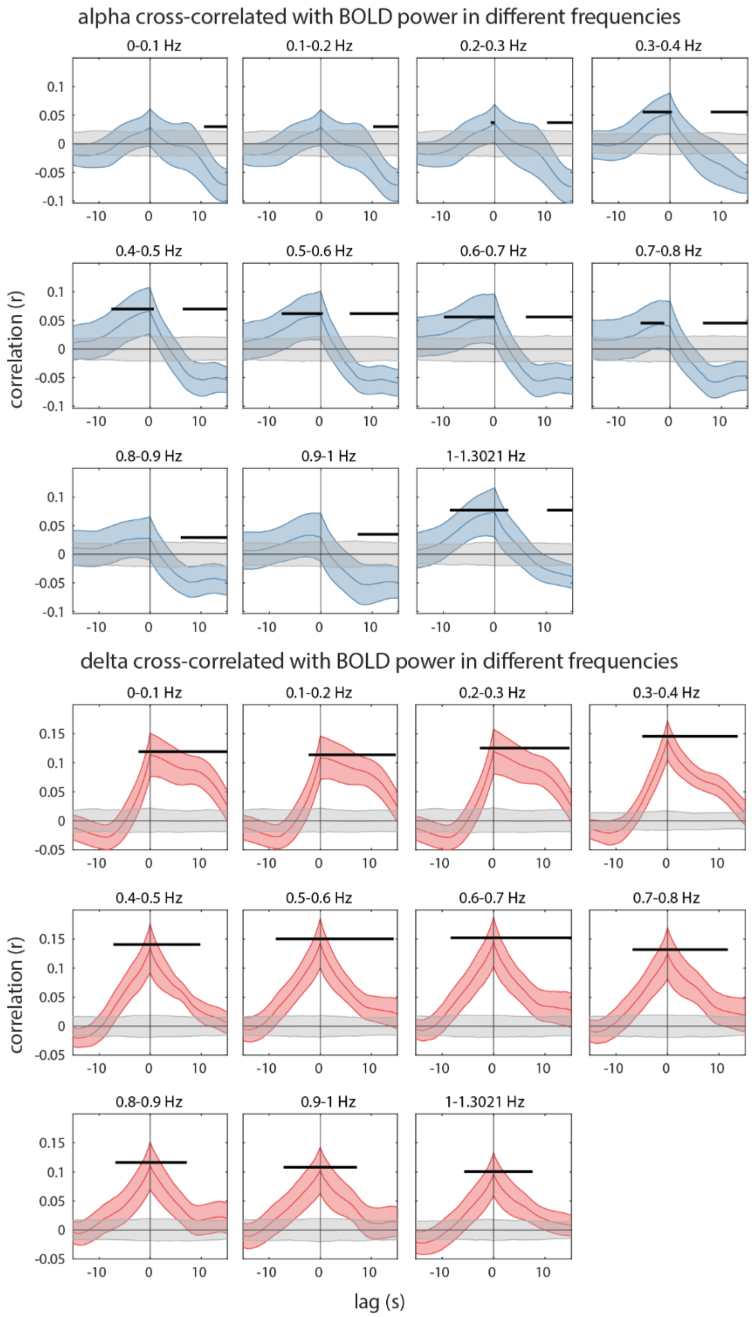


**Supplementary Figure 5:** Alpha and delta EEG power cross-correlated with global cortical BOLD power in different frequencies. Global BOLD signal was calculated from the average of all cortical voxels. Means with 95% CI. Gray lines were obtained from a randomly shifted control, and black bars indicate significant differences (see Methods for details). N=24 (alpha), N=27 (delta).

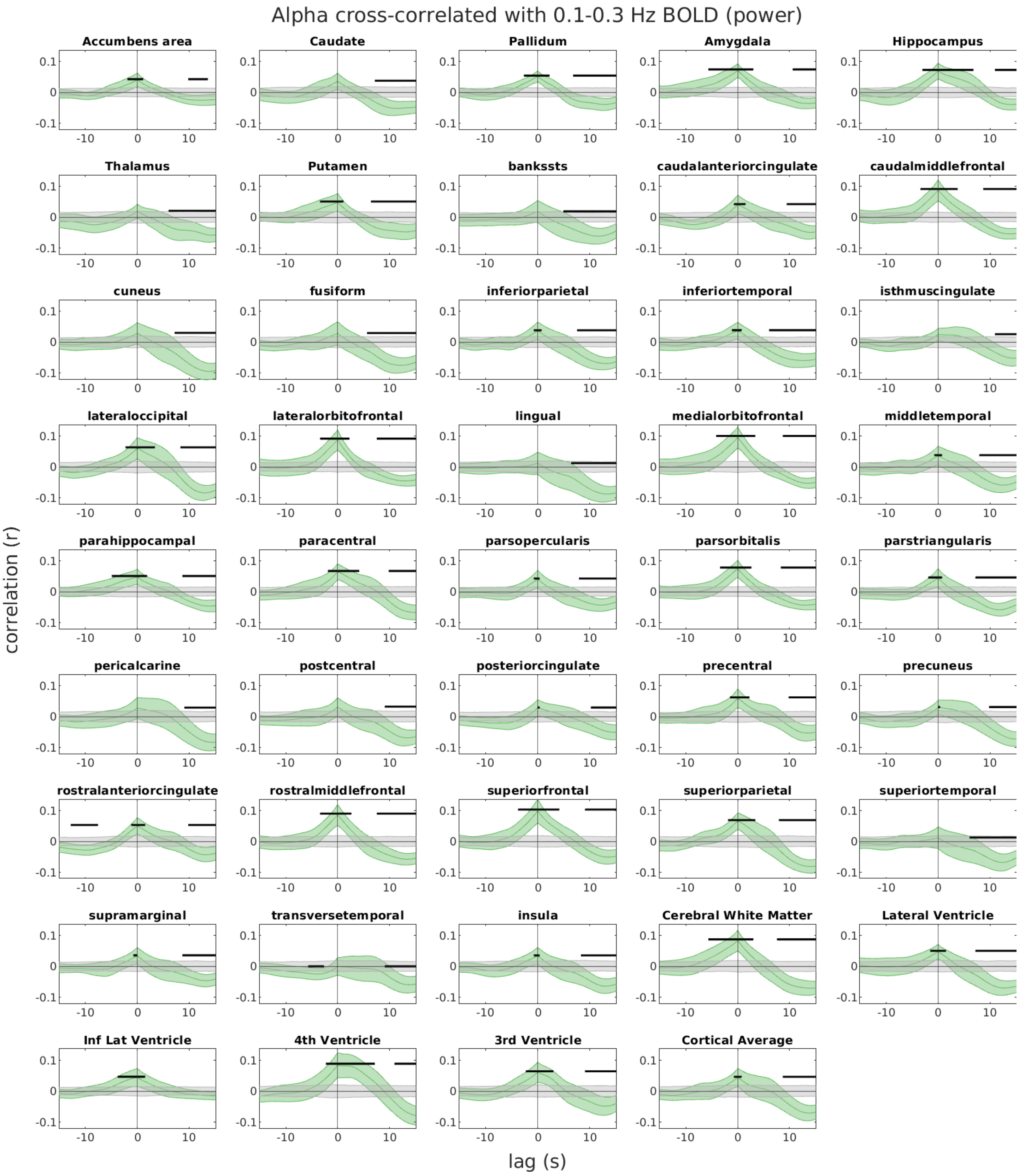


**Supplementary Figure 6:** Alpha power is significantly correlated with fMRI power in the 0.1-0.3Hz range in all regions, with similar patterns to the 0.3Hz and above range. Green lines show group means with 95% CI. Gray lines were obtained from a randomly shifted control, and black bars indicate significant differences (see Methods for details). N=24.

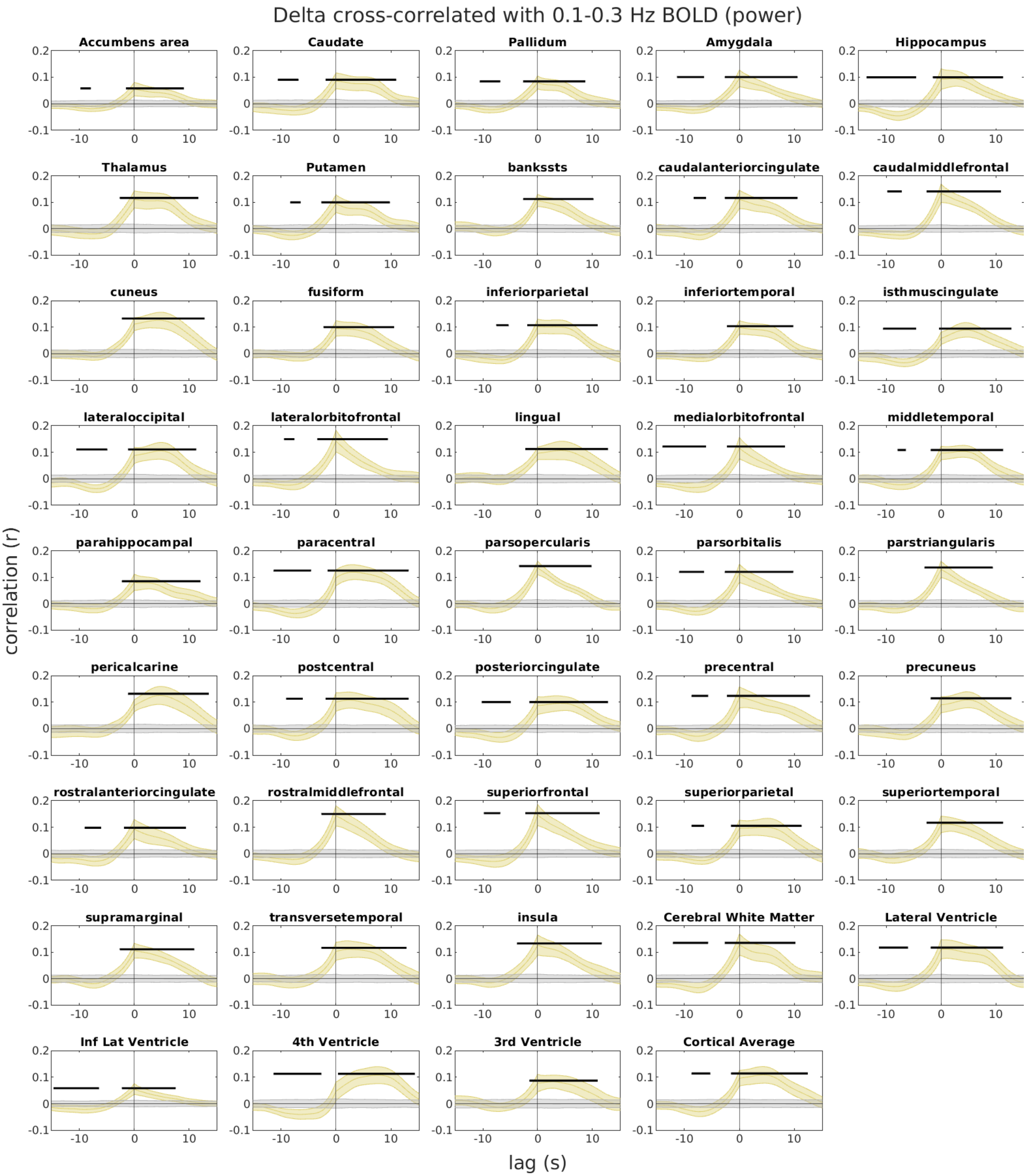


**Supplementary Figure 7:** Delta power is significantly correlated with fMRI power in the 0.1-0.3Hz range in all regions, with similar patterns to the 0.3Hz and above range. Yellow lines show group means with 95% CI. Gray lines were obtained from a randomly shifted control, and black bars indicate significant differences (see Methods for details). N=27.

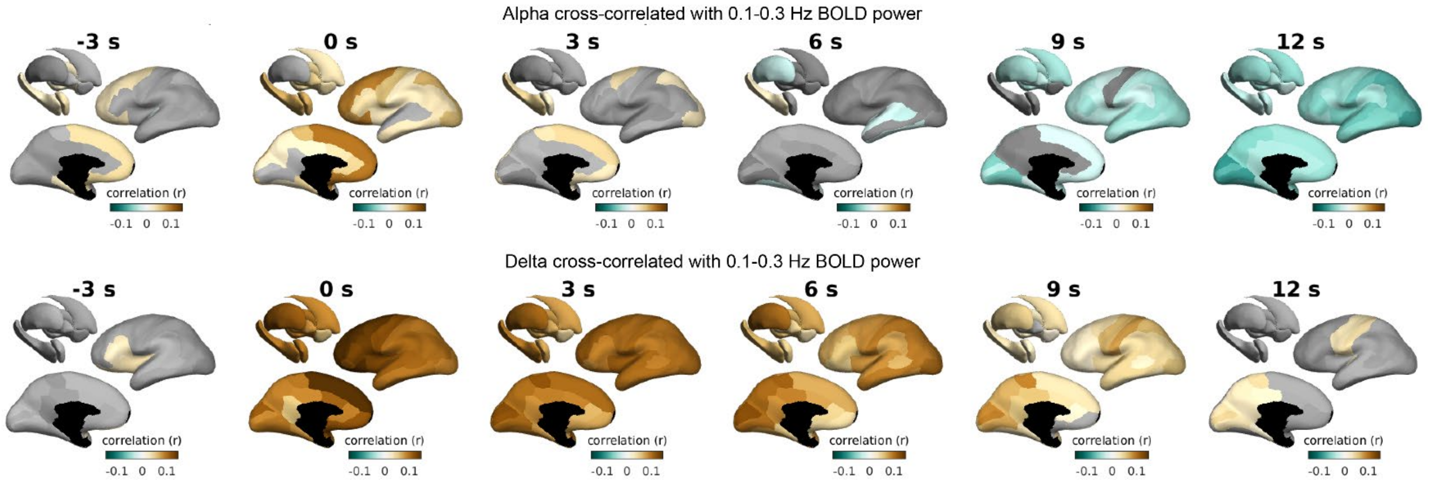


**Supplementary Figure 8:** Alpha (N=24) and delta (N=27) cross-correlations with fMRI power in the 0.1-0.3Hz range displayed on cortical and subcortical surfaces at different lag timings. Non-significant regions are shown in grayscale.

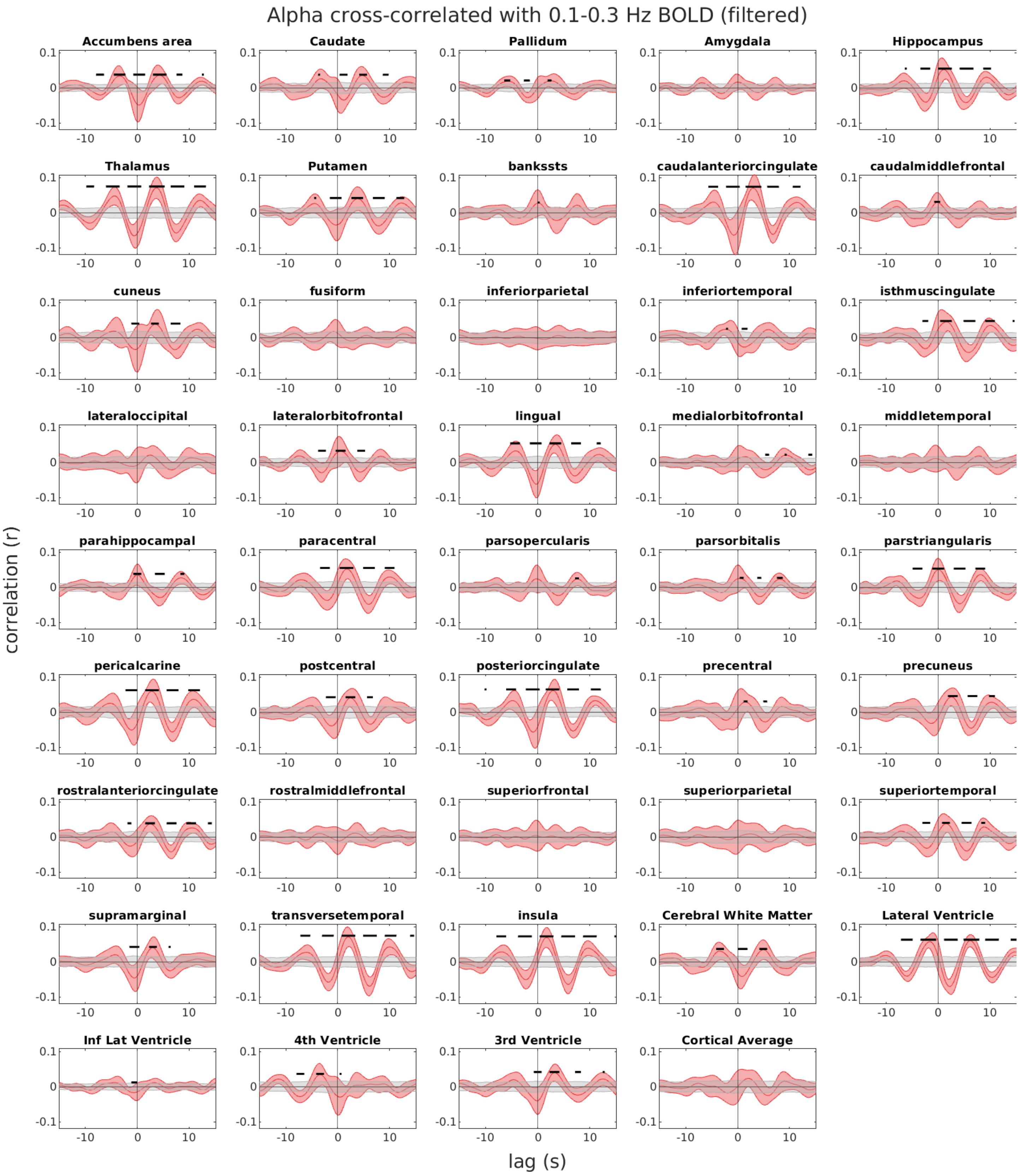


**Supplementary Figure 9:** Alpha power is significantly correlated with the fMRI signal band-passed in the 0.1-0.3Hz range in several regions. Several distinctions are present between these results and the correlations between alpha and fMRI power in the same range; for instance, some regions display band-passed fMRI signals that are strongly correlated with alpha power, but fMRI power in that same range is more weakly related to alpha (e.g. transverse temporal cortex). Conversely, some regions displayed strong correlations between fMRI power in this range and alpha, but no significant correlations when the fMRI signal in this range was not converted to power (e.g. fusiform cortex). Red lines show group means with 95% CI. Gray lines were obtained from a randomly shifted control, and black bars indicate significant differences (see Methods for details). N=24.

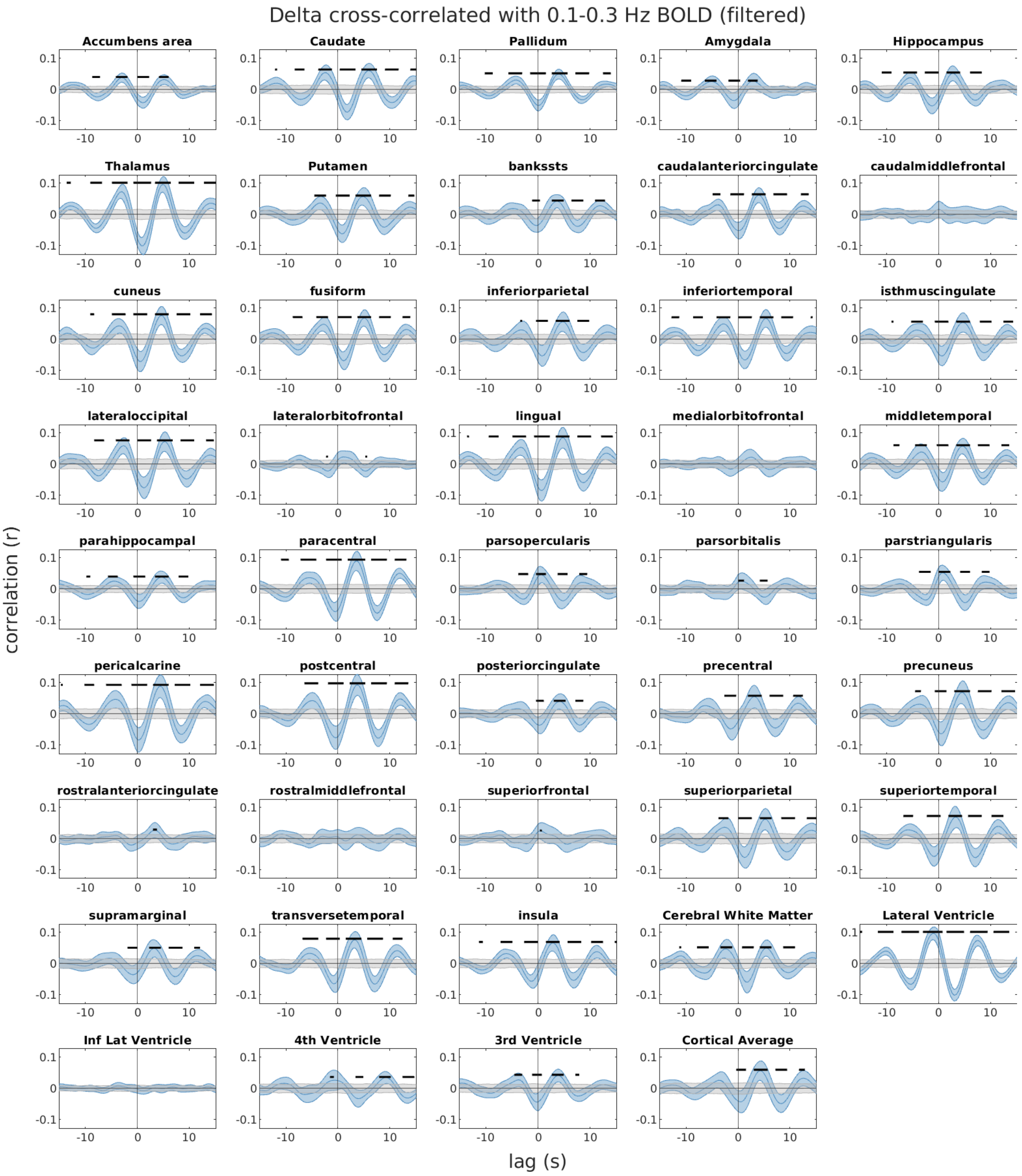


**Supplementary Figure 10:** Delta power is significantly correlated with the fMRI signal band-passed in the 0.1-0.3Hz range in most regions. Patterns differ from equivalent alpha results in lag timings and correlation strengths. Blue lines show group means with 95% CI. Gray lines were obtained from a randomly shifted control, and black bars indicate significant differences (see Methods for details). N=27.

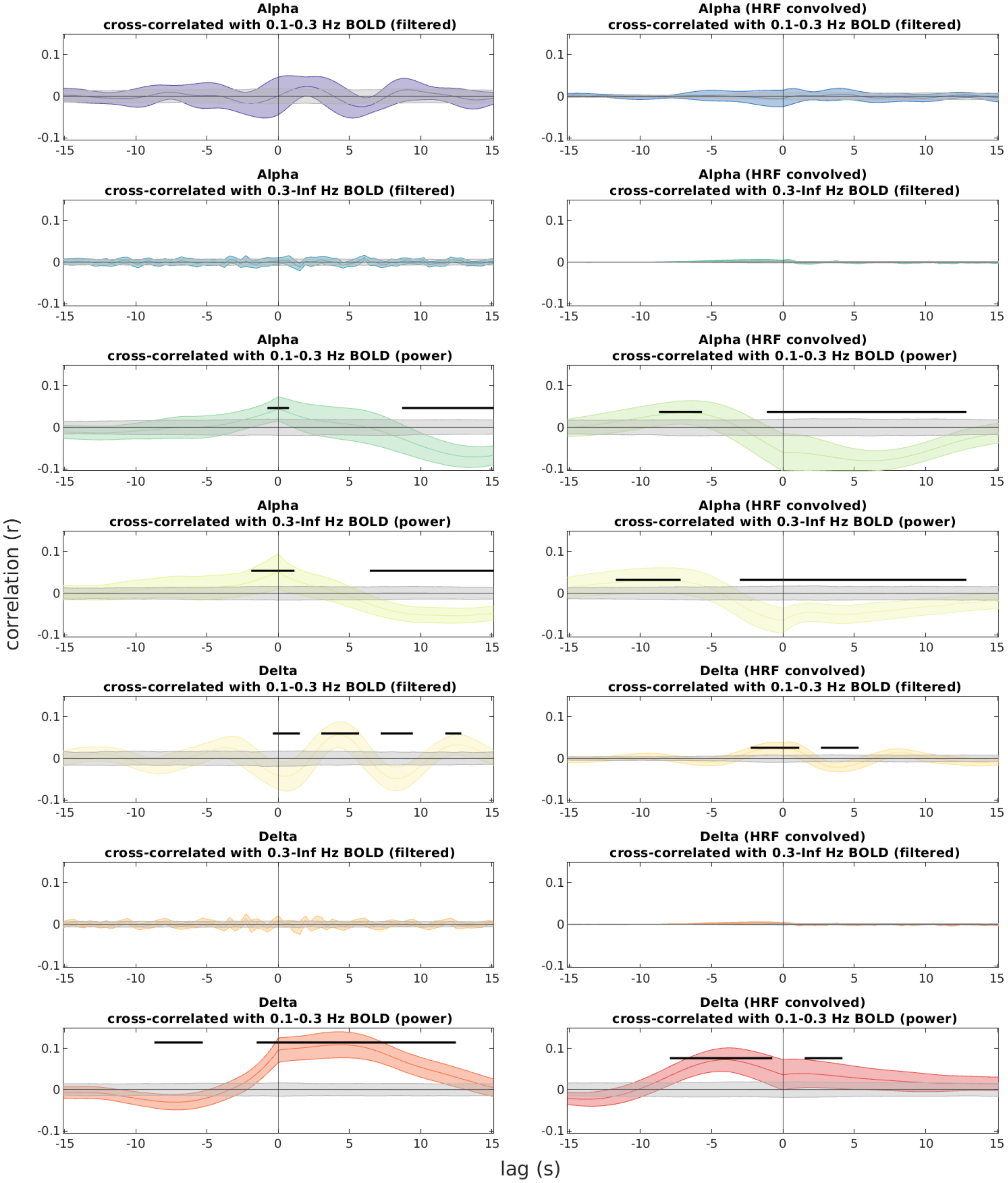


**Supplementary Figure 11:** Convolving the EEG power series with the canonical hemodynamic response function (HRF) did not improve correlation strengths with the cortical average signal. Similar results were obtained in other brain regions. Color lines show group means with 95% CI. Gray lines were obtained from a randomly shifted control, and black bars indicate significant differences (see Methods for details). N=24 (alpha), N=27 (delta).

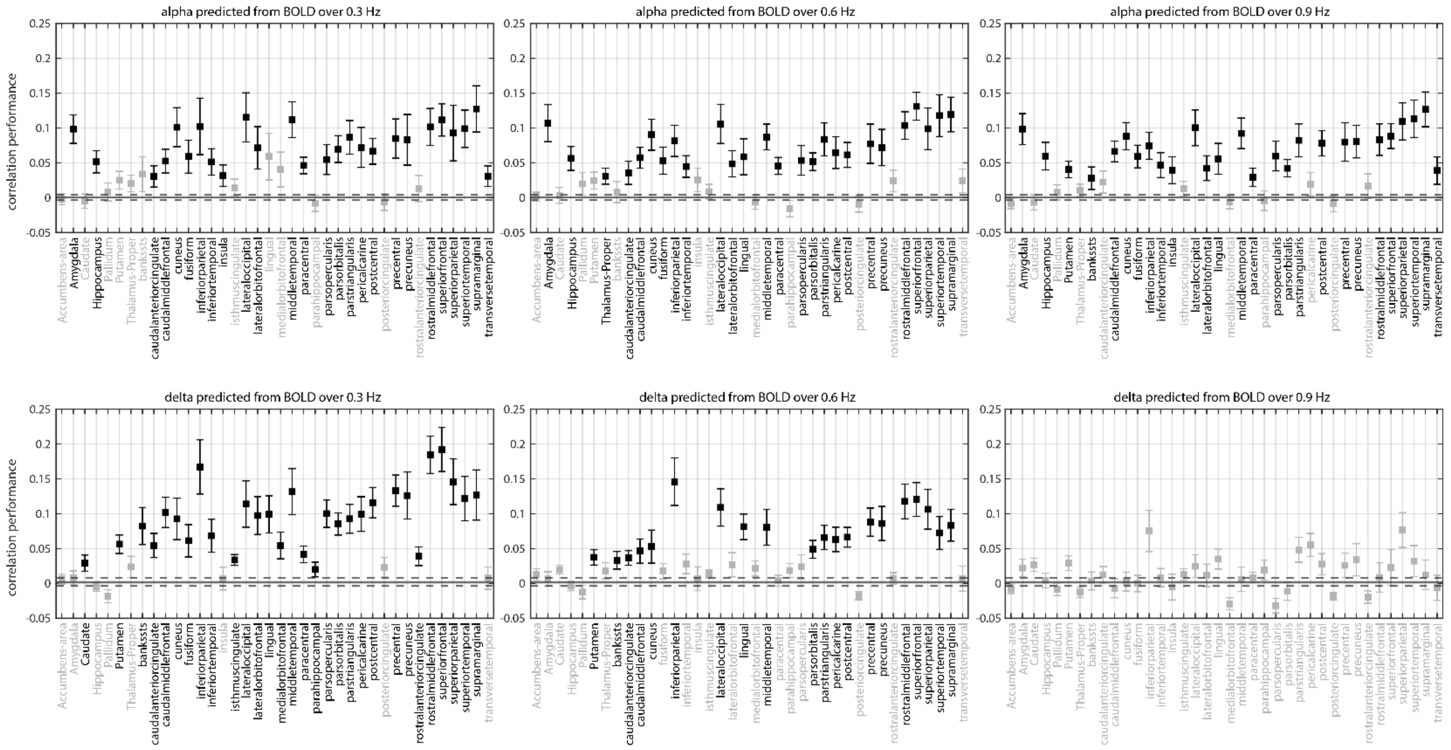


**Supplementary Figure 12:** High-frequency BOLD signals from individual brain regions can predict alpha and delta rhythms. We trained models to predict alpha and delta from one anatomically-parcellated bilateral brain region at a time, with its fMRI activity high-pass filtered using three cutoffs. Regions plotted in black were significantly better than control (paired t-tests, p<0.05, Benjamini-Hochberg correction; control consisted of fMRI data randomly shifted in relation to the EEG). Mean of the control condition is shown as a solid line, with SEM shown as dashed lines. N=24 (alpha), N=27 (delta).

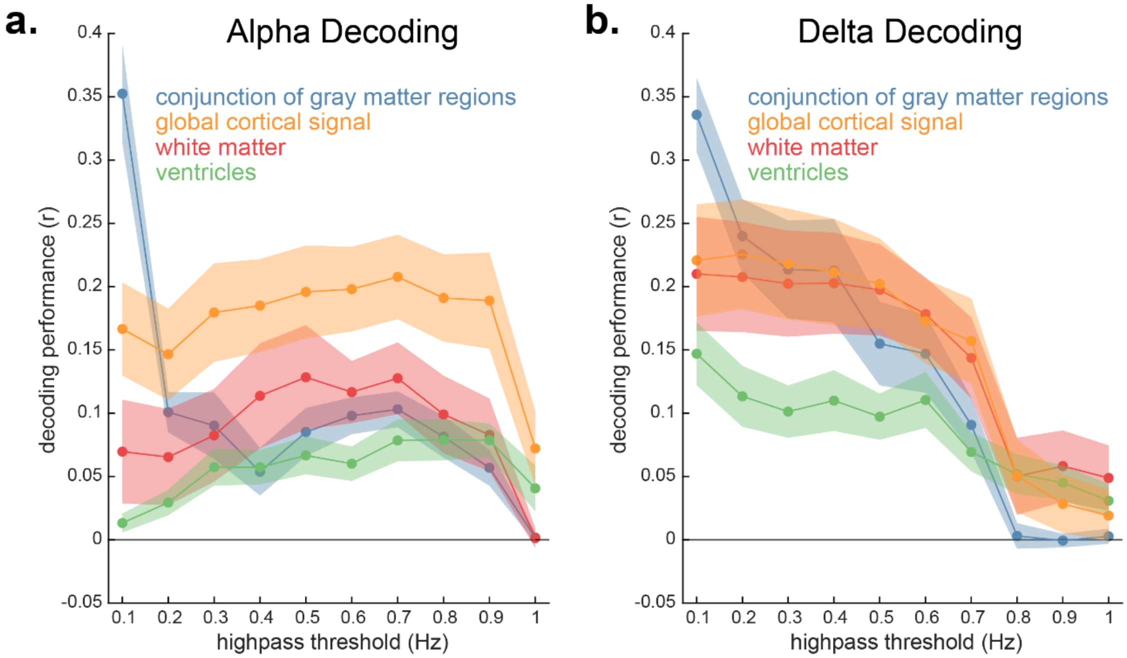


**Supplementary Figure 13:** Temporal profile of decoded high-frequency information across fMRI anatomical regions. ***a–b.*** Brainwide fMRI regions were split into four anatomical conditions: all gray matter regions across the cortex and subcortex (jointly presented to the model), the global cortical signal (average of all cortical voxels), the white matter, and the ventricles (left and right lateral and inferior lateral ventricles, the third ventricle, and the fourth ventricle). These were further split into temporal conditions in which fMRI signals were high-pass filtered progressively higher. Each condition was then used to train models to decode alpha and delta EEG power. Decoding performance consisted of correlation between predictions on held-out subjects and ground truth. ***a.*** Alpha information was decoded from local gray matter signals up to 0.2 Hz (blue lines) and the global cortical signal up to 1 Hz (yellow lines). Decoding from white matter and ventricles was poor. *N*=24. Means with SEM. ***b.*** Delta information was decoded from local and global gray matter signals, along with white matter signals, up to 0.8 Hz (blue and yellow lines). Decoding from ventricles was poor, but superior than alpha decoding. *N*=27. Means with SEM.

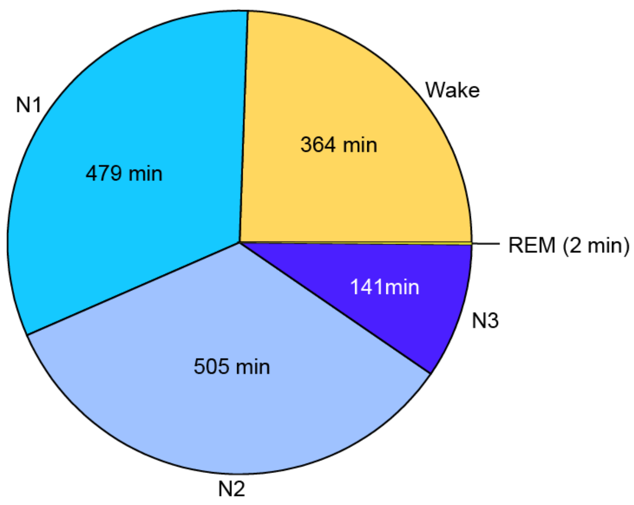


**Supplementary Figure 14:** Total time spent in each sleep stage across all subjects (n=27).
